# Antigenic cartography using hamster sera identifies SARS-CoV-2 JN.1 evasion seen in human XBB.1.5 booster sera

**DOI:** 10.1101/2024.04.05.588359

**Authors:** Wei Wang, Gitanjali L. Bhushan, Stephanie Paz, Charles B. Stauft, Prabhu Selvaraj, Emilie Goguet, Kimberly A. Bishop-Lilly, Rahul Subramanian, Russell Vassell, Sabrina Lusvarghi, Yu Cong, Brian Agan, Stephanie A. Richard, Nusrat J. Epsi, Anthony Fries, Christian K. Fung, Matthew A. Conte, Michael R. Holbrook, Tony T. Wang, Timothy H. Burgess, Edward Mitre, Simon D. Pollett, Leah C. Katzelnick, Carol D. Weiss

## Abstract

Antigenic assessments of SARS-CoV-2 variants inform decisions to update COVID-19 vaccines. Primary infection sera are often used for assessments, but such sera are rare due to population immunity from SARS-CoV-2 infections and COVID-19 vaccinations. Here, we show that neutralization titers and breadth of matched human and hamster pre-Omicron variant primary infection sera correlate well and generate similar antigenic maps. The hamster antigenic map shows modest antigenic drift among XBB sub-lineage variants, with JN.1 and BA.4/BA.5 variants within the XBB cluster, but with five to six-fold antigenic differences between these variants and XBB.1.5. Compared to sera following only ancestral or bivalent COVID-19 vaccinations, or with post-vaccination infections, XBB.1.5 booster sera had the broadest neutralization against XBB sub-lineage variants, although a five-fold titer difference was still observed between JN.1 and XBB.1.5 variants. These findings suggest that antibody coverage of antigenically divergent JN.1 could be improved with a matched vaccine antigen.

## Introduction

New SARS-CoV-2 variants continue to cause significant morbidity and mortality particularly among vulnerable groups despite increasing population immunity to SARS-CoV-2 from infections and vaccinations^1^. COVID-19 vaccines remain an important tool for protecting against infection, severe illness, and Long COVID^2-5^, but vaccine antigen updates may be needed for new variants^6^. Neutralizing antibody titers help assess whether new variants evade vaccine-induced immunity and have been shown to correlate with protection against COVID-19^7-9^.

Antigenic cartography, initially introduced as a new analysis tool to aide interpretation of antigenic differences among influenza strains, uses antibody titers to visualize antigenic differences among strains^10^. Convalescent sera from a first infection (primary infection sera) are often used to generate maps that discriminate viruses based on recognition by antibodies. However, human primary infection sera of new SARS-CoV-2 variants are now rare due to population immunity from prior SARS-CoV-2 infections, vaccinations, or both. Syrian golden hamsters are among the preferred animal models for studying SARS-CoV-2^11-13^ and have been used for generating primary infection sera for antigenic analyses^14-16^. Yet, data directly comparing the similarity between hamster and human antibody responses to the variants are limited^17^, especially to the most recent Omicron variants.

Landscape analyses provide a method for visualizing how complex sera from multiple antigenic exposures from infections and vaccinations recognize different variants of a pathogen^18^. Antibody titers against different viruses are modeled as a landscape over the primary sera antigenic map that defines the antigenic space. Landscapes may allow interpolation of complex serum titers against variants in the map for which direct measurement have not been made. Landscape analyses have previously been included in discussions regarding vaccine updates^19^. However, prior antigenic exposures can shape subsequent responses to vaccines or infections, so complex and primary infection sera may recognize some variants differently^20,21^.

In this study, we generated a unique dataset of neutralization titers against a large panel of SARS-CoV-2 variants, including pre-Omicron and Omicron variants through JN.1, using hamster sera from primary, sequence-confirmed variant infections. We applied antigenic cartography analyses to this serum set to characterize antigenic drift among recent SARS-CoV-2 variants. We further found good comparability between our antigenic map made with hamster sera and our prior antigenic map made with human sera, suggesting that hamster primary infection sera may be valuable for assessing antigenic differences among future variants. Our analyses of complex sera from persons with well-documented, COVID-19 vaccinations and infection exposures also suggests that antigen-matched vaccine boosters could improve protection against the most recent SARS-CoV-2 variants.

## Results

### Hamster primary infection sera exhibit poor cross-neutralization between pre-Omicron and Omicron variants

To assess the suitability of hamster primary infection sera for antigenic analyses to aid selection of antigens for COVID-19 vaccines, we first determined neutralization patterns against a large panel of pre-Omicron and Omicron variants (Fig. 1). Variants tested for pseudovirus neutralization ranged from ancestral to JN.1 variants, but the number of variants tested for each primary infection serum sample was limited by available serum. For pre-Omicron variant infections, neutralization titers were generally highest against the infecting variant, with variable cross-neutralization among pre-Omicron variants and much less cross-neutralization against Omicron variants (Fig. 1A). Infection by the Delta (B.1.617.2) variant generated the highest titers, possibly related to a more robust infection by this variant^22-25^. At the same time, the Delta variant appeared to be more sensitive to cross-neutralization than other pre-Omicron variants, possibly due to better exposure of neutralizing determinants. Omicron sera also showed the highest titers against the infecting variant, with modest cross-neutralization against other Omicron variants and little or no cross-neutralization to pre-Omicron variants (Fig. 1B). Notably, BA.1 and BA2 sera showed no cross-neutralization against JN.1 and XBB-lineage variants. In contrast, BA.5 sera showed cross-neutralization against the JN.1 and XBB-lineage variants, though titers were 24-75-fold lower than against the infecting BA.5 variant. XBB-lineage sera showed cross-neutralization against JN.1 and BA.4/BA.5. JN.1 sera also showed cross-neutralization against BA.4/BA.5 and XBB-lineage variants in a few animals. XBB.1.5 sera showed high neutralization titers against the XBB-lineage variants. The overall lower neutralization titers of BA.1, BA.2, XBB, EG.5.1 and JN.1 sera compared to BA.5, XBB.1.5 and XBB.1.16 sera may reflect variant differences in virus replication in hamsters or differences in immunogenicity and exposure of neutralizing epitopes.

**Figure 1.**
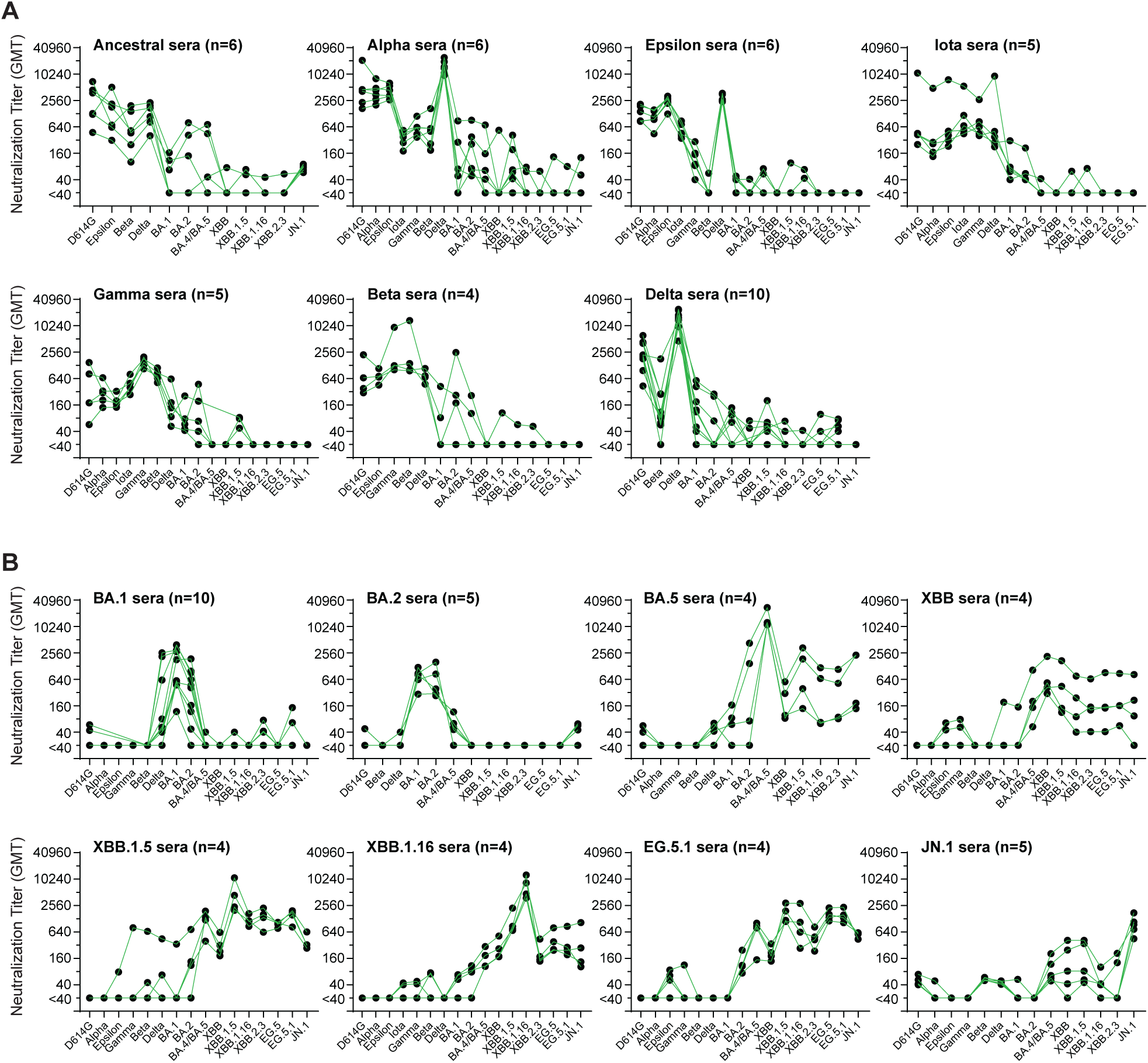
Neutralization titers and specificity against SARS-CoV-2 variants by hamster primary infection sera. (**A**) Hamster pre-Omicron primary infection sera. (**B**) Hamster Omicron primary infection sera. Neutralization titers against the indicated variants in hamster primary infection sera are plotted. Dots indicate the results from individual samples that are connected among titers against different variants. Neutralizations were measured in lentiviral-based pseudovirus neutralization assays. n: sample number.

### Human and hamster neutralization titers correlate well

We next compared neutralization titers from the hamster primary infection sera to the human primary infection sera that we previously reported for pre-Omicron variants and BA.1 variants^26^. A comparison of spike amino acid sequences among primary human infection virus, hamster infection virus, and pseudovirus within given lineages revealed few differences (Fig. S1). The number of differences within a lineage ranged from zero (B.1 in human infection versus pseudovirus) to six (BA.1 in human infection versus hamster infection) spike amino acid differences. In the latter case, most spike substitutions are outside of the spike receptor binding domain. Using the same pseudovirus neutralization assay, we found that human and hamster primary infection sera gave remarkably similar neutralization titers and specificity against the same SARS-CoV-2 variants (Figs. 2 and S2). Overall, both human and hamster neutralization titers were typically highest against infecting variant, with modest cross-neutralization by pre-Omicron primary sera to pre-Omicron variants but comparatively little cross-neutralization to Omicron variants. Likewise, the Omicron primary infection sera did not cross-neutralize pre-Omicron variants.

**Figure 2.**
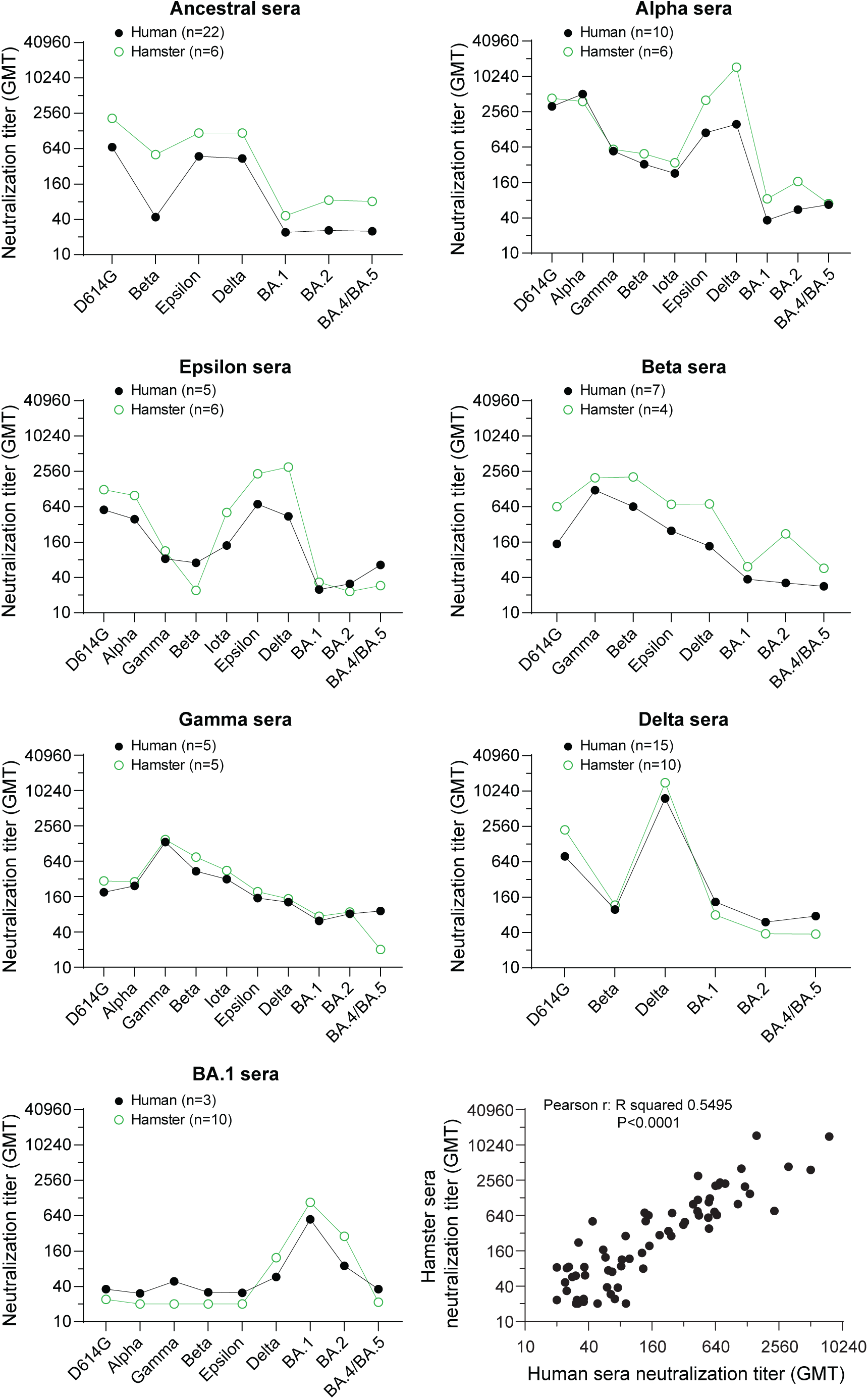
Neutralization titers and specificity against SARS-CoV-2 variants correlate well between human and hamster primary infection sera. Neutralization titers (GMT) against the indicated variants in human and hamster primary infection sera of the indicated SARS-CoV-2 variants infection are plotted. A Pearson correlation among all neutralization titers (GMT) of all human and hamster sera is analyzed. Dots indicate the GMT. n: sample number. All data of human primary infection sera were reported previously^26^. Neutralization titers of the individual serum samples against the variants are shown in Supplemental Figure S2. Spike genotypes of the hamster infections and pseudoviruses used for neutralizations are listed in Supplemental Table S4. Spike genotypes of the human infections are listed in Supplemental Tables S7 and S8. A full comparison of the genotypes of all spikes is shown in Supplemental Figure S1.

### Antigenic maps created from human and hamster primary infection sera closely resemble each other

We applied antigenic cartography to our extended hamster sera neutralization dataset, which allowed us to measure antigenic relationships to and among these newer Omicron variants that we could not measure using human primary infection sera alone. As reported previously for primary human sera maps ^26-28^, Alpha, Delta, Epsilon, Beta, and Gamma variants formed a distinct antigenic cluster from the early Omicron variants BA.1, BA.2, BA.2.12.1, and BA.4/BA.5 (Fig. 3A, Table S1). These variants have similar antigenic relationships to one another on a map made only with hamster sera (Fig. 3B, Procrustes arrows point from the position in one map to another in Fig. 3C), indicating the potential suitability of hamster primary infection as a substitute for primary human infection sera. Distances from D614G to each of the variants is also similar between maps, with the greatest differences seen for BA.1 (1.3 antigenic units, AU) and BA.4/BA.5 (0.7 AU) (Table S1). On the hamster map, the newly emerged XBB-lineage variants form an additional cluster (XBB cluster) furthest away from the ancestral variants and distinct from BA.1 and BA.2 (Fig. 3B). With this addition of XBB-lineage variants in the hamster map, BA.4/BA.5 moves away from BA.1 and towards the more recent XBB-lineage variants (Fig. 3B). Notably, JN.1 also clusters with BA.4/BA.5.

**Figure 3.**
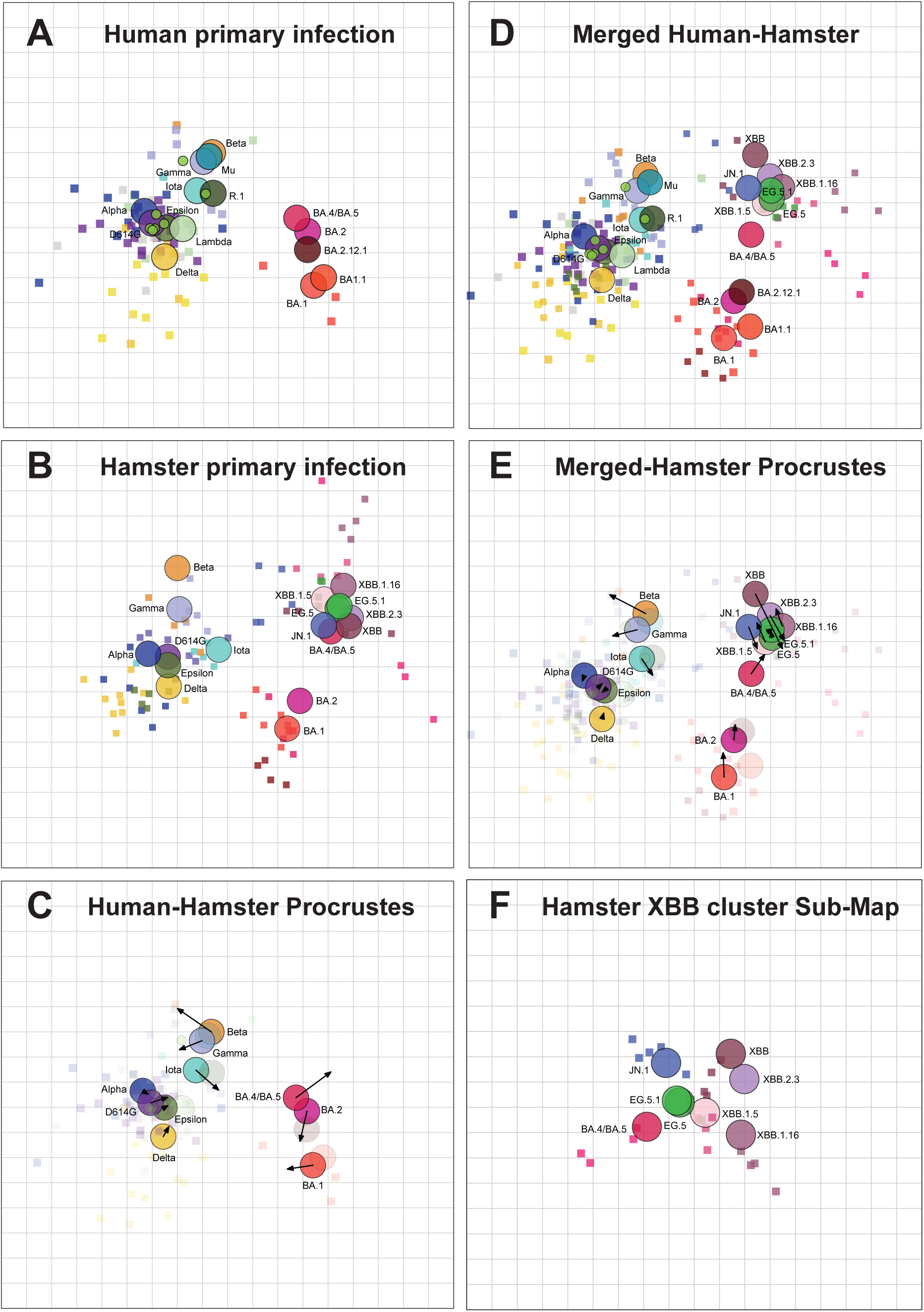
Antigenic maps made with neutralization titers from human and hamster primary infection sera closely resemble one another. Antigenic maps were made using antigenic cartography with titers for primary infection sera from **(A)** Humans and (**B**) Hamsters. (**C**) Comparison of virus positions between human primary infection map and hamster primary infection map. Arrows point to virus positions from human map to hamster map, **(D)** Merged human-hamster map containing primary infection sera from both humans and hamsters. (**E**) Merged human-hamster map compared to the hamster primary infection map. (**F**) Submap including only the hamster XBB cluster. Each grid-square side corresponds to a 2-fold dilution in the pseudovirus neutralization assay. Antigenic distance is measured in any direction on the grid. Antigens are shown as circles and labeled. Sera are shown as squares and are colored by infecting variant. The human primary infection map was reported previously ^26^.

Given that the human and hamster sera measured similar antigenic relationships among the viruses, we created a merged human and hamster map including all data from both datasets (Fig. 3D). The merged antigenic map had similar error and fit to titer data as the individual human and hamster primary infection map (Fig. S3A) and had robustness in the positioning of viruses and sera when adding titer and antigen noise, as well as when resampling the data (Fig. S4). The merged map shows similar antigenic relationships to the hamster map (Fig. 3E), with the greatest shifts in position in the XBB cluster. Compared to the hamster map, JN.1 and BA.4/BA.5 diverge in the merged human-hamster map, with JN.1 remaining with the XBB cluster and BA.4/BA.5 assuming a position between the XBB cluster and the early Omicron cluster (Fig. 3E).

Based on dimensionality testing, we observed that the titers were better fit on maps made in three dimensions (Fig. S3C and S3D). When we evaluated differences in antigenic positions between the 2D and 3D merged maps, we observed some of the most substantial shifts were within the XBB cluster, suggesting that the antigenic relationships in the 2D map were not as accurately representing the underlying data in this region of the map (Fig. S5A). We thus evaluated whether antigenic relationships among the variants in the XBB cluster differed if evaluated only using hamster sera raised against these variants. Both the serum and antigen positions shifted, in some cases substantially, between the full map and the XBB cluster map (Fig. 3F). This shifting of antigenic positions within a cluster was not observed when the same analysis was performed on the earlier variant cluster (Fig. S5B). Notably, distances were greater between JN.1, XBB.1.5, and BA.5 on the XBB cluster map (5 to 6-fold difference between each antigen and each other) compared to the full map (1 to 2.5-fold difference) (Fig. 3B and 3F). This suggests that the primary infection sera against XBB variants saw differences among these antigens that sera raised against early variants and Omicron were not able to detect, and the full map, which must accommodate all titers, did not fit the titers in the XBB cluster as well.

### Multiple antigenic exposures increase neutralization breadth against the variants

We next investigated neutralization breadth in sera from persons who had one of the following multiple antigenic exposures (Fig. 4A): 1) three doses of the ancestral mRNA COVID-19 vaccine (V3)^26^; 2) three doses of the ancestral mRNA COVID-19 vaccine and a post-vaccination infection (PVI) during BA.1 outbreak (V3+PVI); 3) four doses of the ancestral mRNA COVID-19 vaccine (V4); 4) three doses of the ancestral mRNA COVID-19 vaccine and one dose of the bivalent (ancestral+BA.5) mRNA COVID-19 vaccine (V3+Bi); 5) four doses of the ancestral mRNA COVID-19 vaccine and one bivalent mRNA COVID-19 vaccine (V4+Bi)^29^; and 6) three or four doses of the ancestral mRNA COVID-19 vaccine, with or without a PVI before or after one bivalent mRNA COVID-19 vaccine followed by one XBB.1.5 mRNA vaccine (V3 or V4+bivalent+XBB.1.5 with or without PVI, referred to as XBB.1.5 booster group*). In the XBB.1.5 booster group*, six individuals had no reported PVI. Based on time of infection, the PVIs in the XBB.1.5 group* included three presumed BA.1 PVIs, four presumed BA.5 PVIs after three doses of the ancestral COVID-19 vaccine, and three presumed XBB PVIs after the bivalent booster (Table S2). Variant assignments of PVIs were based on the dominant circulating variant at the time of the documented infection.

**Figure 4.**
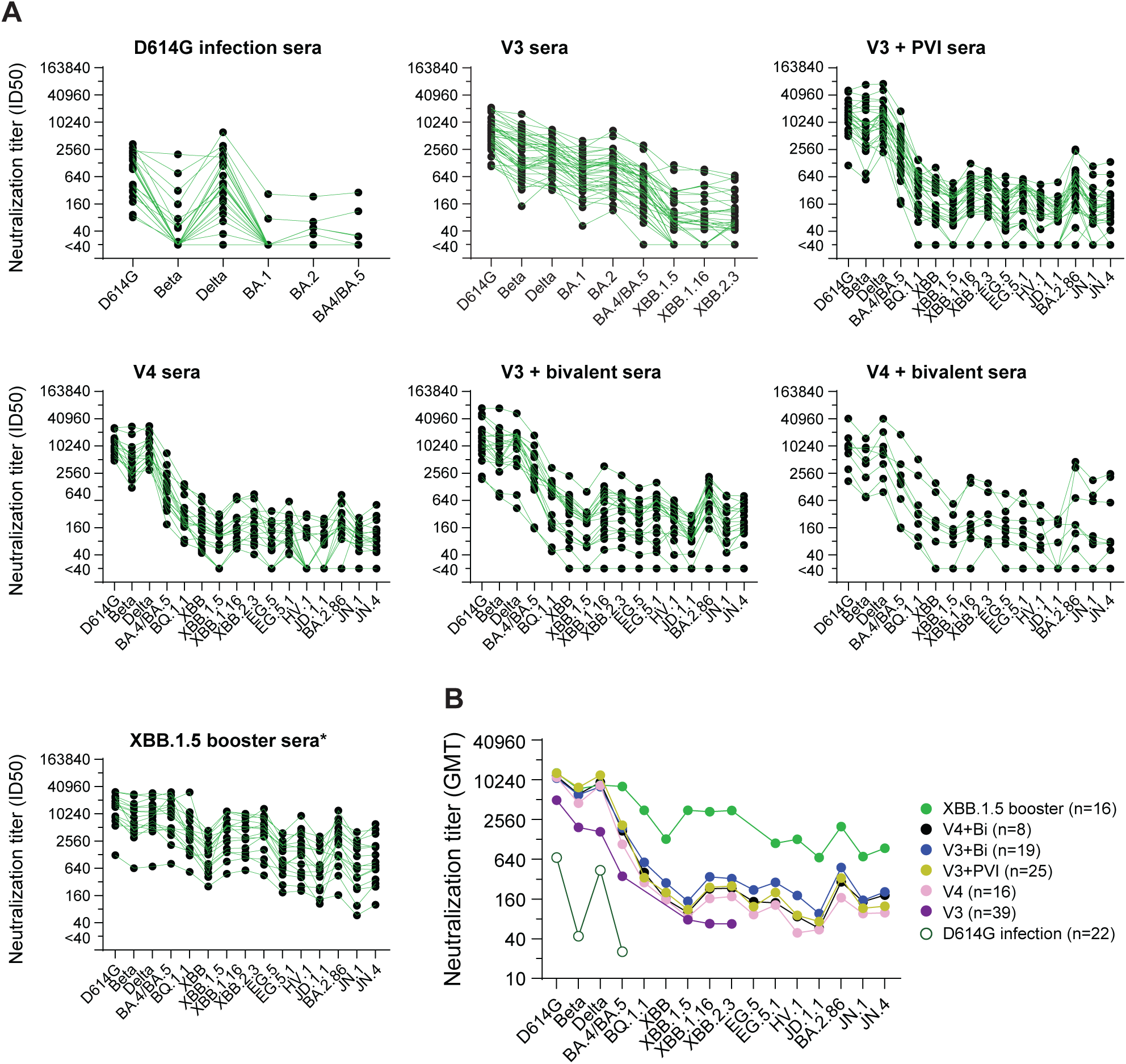
Neutralization titers and specificity against SARS-CoV-2 variants in human sera after multiple antigen exposures. (**A**) Neutralization titers represented as 50% inhibitory dilutions (ID_50_) against pseudoviruses bearing spike proteins from the indicated variants are plotted for the human sera after multiple antigen exposures. Dots indicate the results from individual samples that are connected among titers against different variants. (**B**) The comparison of neutralization titers (GMT) against the indicated variants in each sera panel. n: sample number. All data of human primary infection sera were reported previously^26^. All data of V3 sera, except neutralizations against the stains of XBB-lineages, were reported previously^26^. Neutralization titers against D614G, BA.4/BA.5, BQ.1.1, XBB and XBB1.5 in the sera panels of V4+Bi, V3+Bi, V3+PVI and V4 were reported previously^29^. XBB.1.5 booster group* includes: six individuals without reported PVIs; three individuals with presumed BA.1 PVIs; four individuals with presumed BA.5 PVIs after three doses of the ancestral COVID-19 vaccine; and three individuals with presumed XBB PVIs after the bivalent booster. PVI infections were assigned according to the time of infections.

Overall, all groups that received vaccinations had higher neutralization titers and more breadth against the variants than the group that had only D614G infection (Fig. 4A-B). The different mutations in variant spikes compared to D614G are summarized in Fig. S6A. Groups that had heterologous antigenic exposures from XBB.1.5 or bivalent COVID-19 boosters or a BA.1 PVI after vaccinations with the ancestral COVID-19 vaccine had neutralization titers that trended higher than those from the group that had only the ancestral COVID-19 vaccine (V3) when measured against recently emerged XBB-lineage (XBB, XBB.1.5, XBB.1.16, XBB.2.3, EG5, EG.5.1, JD.1.1 and HV.1) and BA.2.86-lineage (BA.2.86, JN.1 and JN.4) variants (Fig. 4A-B). Neutralization titers against XBB-lineage and BA.2.86-lineage variants were 4-22-fold lower than titers against BA.4/BA.5 after the bivalent COVID-19 vaccine (Fig. 4B and S6B), indicating that XBB-lineage and BA.2.86-lineage variants exhibit substantial immune evasion from immunity elicited by prior vaccines, even after multiple immunizations. Notably, compared to the V4, V3+Bi, V3+PVI, and V4+Bi groups, the XBB.1.5 booster group* had higher neutralization titers against XBB-lineage and recently emerged variants, including JN.1. This finding is consistent with recent reports^30-38^ although the XBB.1.5 booster-induced neutralization titers against JN.1 are slightly different among the studies. The XBB.1.5 booster also boosted titers against BA.4/BA.5 (Fig. 4B and S7B), consistent with other reports showing back-boosting to earlier variants^31,39-41^. However, neutralization titers against JN.1 were five-fold lower than titers against XBB.1.5 (Figs. 4B, S6B, and S7).

### Sera from the XBB.1.5 booster group have flatter antibody landscapes and higher titers to newer variants than sera from groups with other combinations of vaccinations and pre-XBB variant infections

Antibody landscapes of the multi-exposure sera described above were constructed by fitting a cone-shaped landscape to neutralization titer data. The hamster map was used as the base map in the antibody landscapes. All vaccination groups with or without a PVI had antibody titers, and thus detectable landscapes, across variants (Fig. 5A-E). The landscapes were similar to one another, except for the XBB.1.5 booster sera* landscape. The V4 sera had the least breadth to antigenically distinct variants. The V4+Bi sera had slightly increased breadth, followed by the V3+PVI, and V3+Bi sera. Landscapes were high for pre-Omicron and early Omicron variants (BA.4/BA.5) but dropped against XBB-lineage variants and JN.1. In contrast, the XBB.1.5 booster sera* landscape was high across the map, including against all XBB-lineage variants. Similar patterns were observed for landscapes plotted above the XBB cluster hamster map (Fig. S8).

**Figure 5.**
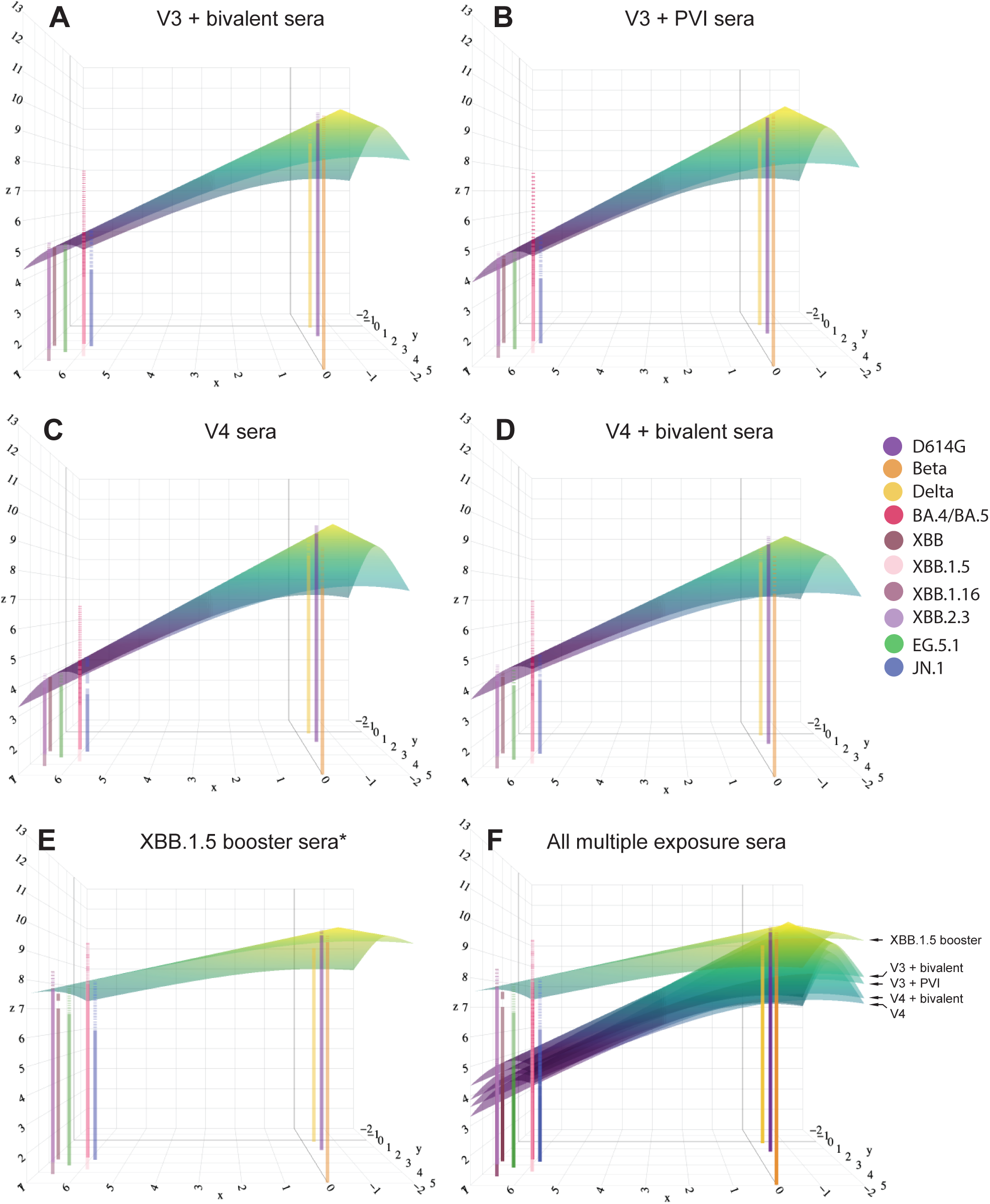
Antibody landscapes of multiple antigen exposure sera groups show that the XBB.1.5 COVID-19 booster vaccine increases magnitude and breadth of antibody titers against newer variants. **(A–E)** Antibody landscapes generated with (**A**) V3+Bi vaccination sera, **(B)** V3+PVI sera, **(C)** V4 vaccination sera, **(D)** V4+Bi vaccination sera, and **(E)** XBB 1.5 booster sera*. (**F**) Each multiple antigen exposure serum group is stacked on top of one another for visualization. All antibody landscapes were plotted above the full hamster antigenic map seen in Figure 3B. The x and y coordinates indicate the location of the variants in the 2D map with measured titer plotted on the z axis in a third dimension above the antigenic map. A cone landscape was fitted and plotted above each 2D map with each landscape being fitted with its own slope. Each landscape is a representation of the average of all individual landscapes within that distinct multiple antigen exposure group. The peak of the cone landscape represents the highest titer measured against any of the antigens. This peak was consistently located above D614G. Lines extending from the x and y axis into the z axis represent the average log GMT against the antigen located at that x,y coordinate. Residuals between measured and predicted titers are represented by a dotted line above or below the landscape corresponding to each antigen.

Antibody titers against Beta, BA.4/BA.5, XBB.1.5 and JN.1 were less well-fit by the cone-shaped landscape as seen by their large residual titers. This was particularly true for BA.4/BA.5, which had measured titers at least two-fold (1 AU) greater than titers predicted from the cone landscape. In contrast, measured titers for JN.1 were more than 1 AU below the titers predicted from the cone landscape (Table S3). This suggests that a cone landscape may not accurately fit titers when plotted above an antigenic map that uses primary infection sera from newly emerged variants such as XBB.1.5. Notably, the large drop in titer for the vaccine sera between BA.4/BA.5 and the XBB variants, despite these viruses being close together on the primary antigenic map, suggests that any landscape or surface may not fit these data well and that primary infection sera may recognize epitopes on the variants differently from multi-exposure sera.

### Repeat boosting decreases the antigenic distance from D614G to BA.5 more than to other variants, even for antigenic histories without BA.5 exposure

Multi-dimensional scaling analyses were also created for multi-antigen exposure sera groups that were described previously^26-28,42-48^. These visualizations cannot be interpreted as antigenic maps but can be useful for visualizing fold-drop in neutralization from infecting strains (*e.g.*, D614G) to other strains. Maps made with multi-antigen exposure sera show that JN.1 and XBB.1.5, as well as other XBB-lineage variants, were more distant from D614G in all serum sets except following XBB1.5 booster vaccination, which decreased antigenic distances to these variants. BA.4/BA.5 was consistently situated closely to D614G (Fig. 6). This contrasted with the hamster and merged human/hamster maps, where JN.1 and BA.5 clustered with the XBB variants, between 6.5-7.9 AU from D614G. One hypothesis is that vaccination with the ancestral variant boosted immunity to a shared epitope on BA.4/BA.5 that is less present on the XBB variants. This difference was not strongly recognized by primary infection sera, potentially because after only a single exposure, the shared epitope between the original variants and BA.4/BA.5 was only partially recognized.

**Figure 6.**
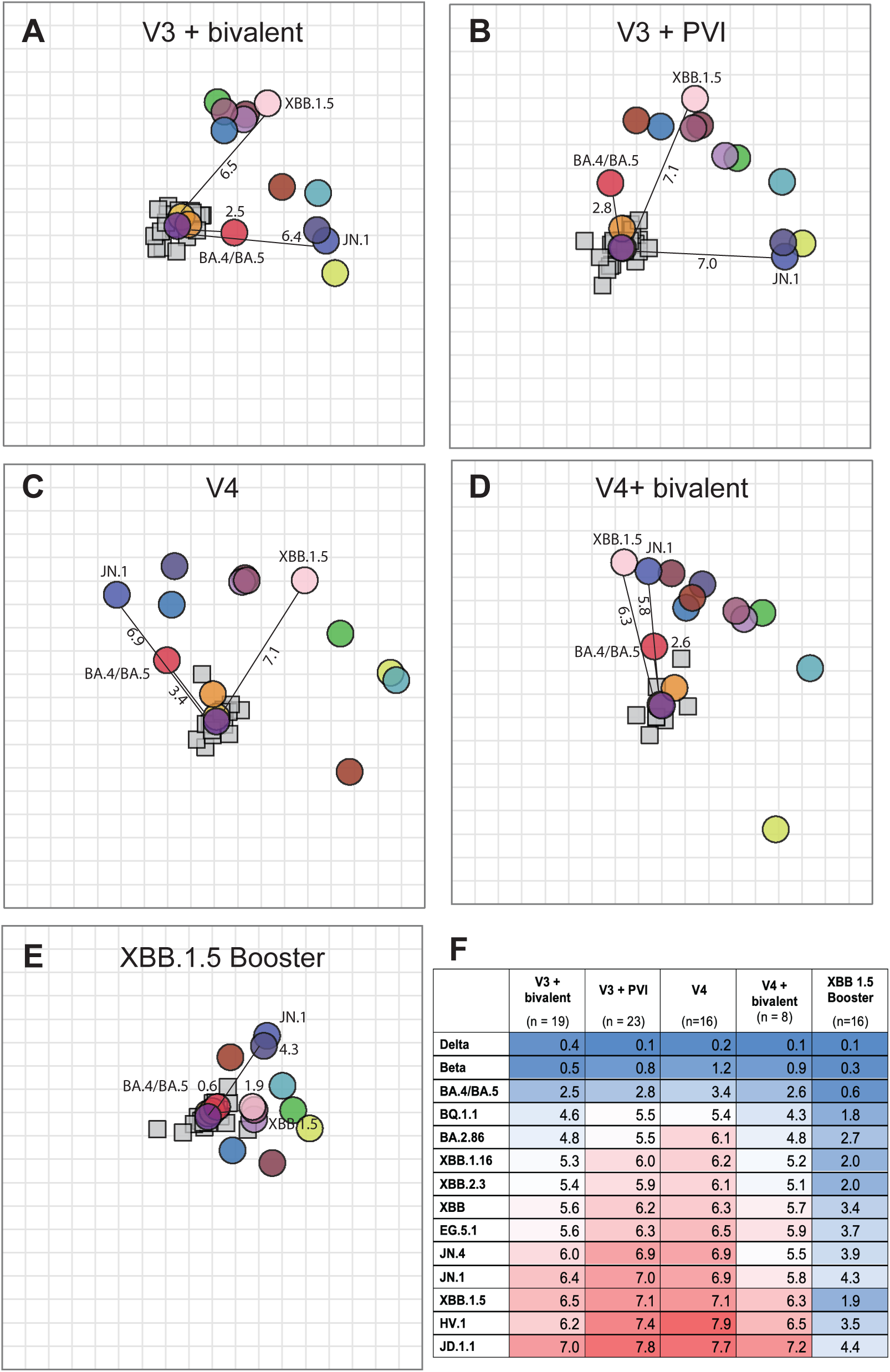
Serum clustering visualizations showing that distances between newer variants and D614G decreases with XBB.1.5 booster. (**A-E**) Antigenic cartography tools were used to measure antigenic distances between D614G and other variants for different multiple antigen exposure groups. (**A**) V3+Bi vaccination sera, **(B)** V3+PVI sera, **(C)** V4 vaccination sera, **(D)** V4+Bi vaccination sera, and **(E)** XBB.1.5 booster sera*. Lines indicate antigenic distances from ancestral D614G to newer variant. Sera are shown as small squares and viruses as colored circles. **(F)** Antigenic distances between D614G and newer variants for each multiple antigen exposure group.

## Discussion

In this study, we extend prior studies by demonstrating that human and hamster primary infection serum titers correlate well. We also provide an extensive primary serum antigenic map that includes ancestral to JN.1 variants, which we discuss in relation to other published antigenic maps and human sera following multiple antigenic exposures. Finally, we show that those who received an XBB.1.5 booster had large increases in antibody titers against newly emerged variants. The combined antigenic and neutralization titer analyses provide insights into variant evolution and contribute to the development of a framework for evaluating antigenic differences among emerging SARS-CoV-2 variants to aide decisions about vaccine antigen updates.

Overall, our primary human and hamster maps show comparable orientations and similar topology to other maps created with human and hamster sera^14-17,28,48^. The consistency between human and hamster maps over a two-year span show that hamster sera are a suitable surrogate for creating antigenic maps to surveil SARS-CoV-2 antigenic variation and predict immune escape. Our merged human-hamster primary infection map positions BA.4/BA.5 and JN.1 in a cluster with newer XBB Omicron variants, suggesting shared epitopes. Previous studies have included limited number of samples or no sera raised against newer variants^15-17,26,39,49^. By including additional single exposure hamster sera for BA.1, BA.4/BA.5, XBB variants and JN.1, our study helps to increase resolution in this region of antigenic space.

Interestingly, we found that the full hamster and merged hamster-human antigenic maps did not accurately fit titers for sera and antigens in the XBB cluster. When we made a map using only the XBB variants and sera raised against these viruses, we found that the antigens moved apart (Fig. 3F). The antigenic distances observed in the XBB cluster map better matched the differences observed by XBB1.5 booster sera*. These findings suggest that there may be multiple ways to use antigenic maps to interpret complex serological data. The map including all antigens and sera is useful for tracking antigenic evolution and identifying clusters. However, greater discrimination among variants within a cluster, which could be important for decisions about vaccine updates, may be achieved by making maps using only sera and antigens from that cluster.

Building upon antigenic cartography, antibody landscapes have been useful for characterizing neutralization profiles of individuals who have had multiple antigenic exposures^18,26,28,39,48,50^. The height of an accurate antibody landscape can be used to infer antibody titers against variants that were not used in making of the landscape^18^. It has been common for antibody landscapes generated with multiple ancestral SARS-CoV-2 vaccinations and infection histories to assume a cone landscape, often with a peak at D614G (ancestral strain) with the landscape gradually tapering as antigenic distance from D614G increases^26,28,39,48,50^. Our study found that booster vaccines or PVIs containing Omicron moderately boosted neutralizing antibodies against Omicron variants more than repeated vaccination with ancestral strain, which has been noted elsewhere^50^. In addition, we show that booster vaccination with XBB.1.5 increases neutralizing antibodies and thus the landscape even higher than prior bivalent vaccinations or PVIs. Previous studies have suggested that vaccination with D614G containing vaccines may blunt antibody responses to the new antigens and that repeat antigen exposure may be needed to boost response to the new antigen^51^. Compared to the prior booster vaccination that included two antigens, D614G and BA.4/BA.5, the XBB.1.5 booster only contained one antigen, which, taken with the higher titers, suggests that including the antigen of interest and excluding D614G may improve neutralizing antibody titers against the antigen of interest as well as antigenically related antigens.

Residual titers plotted on the antibody landscape offer a glimpse of how well the landscape can predict titers against a particular variant. In prior antibody landscapes that use antigenic maps with BA.4/BA.5 as the latest variant, the landscapes with plotted residuals show landscapes that accurately fit antibody titers^26,50^. However, in landscapes that include XBB and BQ variants in their antigenic map, the measured titers against BA.4/BA.5 have generally been greater than what the landscape predicts^39^. In our study, titers for BA.4/BA.5 are also consistently higher than the fitted landscape, including in individuals without a history of BA.4/BA.5 infection. This finding could reflect boosting or affinity maturation of cross-neutralizing antibodies to shared epitopes in BA.4/BA.5 or better exposure of such epitopes in BA.4/BA.5 than XBB variants, even for those only receiving the ancestral vaccine.

Our multiple antigenic exposure results are consistent with previous findings of decreased distance between D614G and Omicron variants in sera from individuals boosted with various Omicron variants (BA.1, BA.5 or XBB.1.5)^26,30,42,46^. The visualizations of multi-antigen exposure sera in this study show that bivalent vaccination with the ancestral strain and BA.4/BA.5 variants shortens the antigenic distance between D614G and BA.4/BA.5 to a greater extent than between D614G and XBB.2.3 and XBB.1.16, as expected given that individuals were exposed to both D614G and BA.4/BA.5. However, we also observe that four doses with the ancestral strain also reduces the distance between D614G and BA.4/BA.5 more than D614G and XBB.2.3 and XBB.1.16, which was not expected, because these individuals had not been exposed to BA.4/BA.5. This is a continuation from Wang et. al. that shows decreased distance between D614G and BA.4/BA.5 when ancestral strain vaccination increases from two to three doses^26^. This observation suggests there may be shared antigenic epitopes between D614G and BA.4/BA.5 that are not detectable using primary infection sera but are present following repeat exposure to the ancestral variant due to vaccination. These results, along with differences in variant neutralization by primary infection sera compared to multi-antigen exposure sera, highlight how past antigenic exposures from infections, vaccines, or both can impact subsequent vaccine responses.

Importantly, in sequential exposures to multiple SARS-CoV-2 antigens, the latest new antigen not only elicits titers against the most recently exposed antigen, it also back-boosts titers to previously exposed antigens, as shown here by the boosted titers against BA.4/BA.5 after the XBB.1.5 vaccine booster (Fig. 4B and S7B). Such back-boosting from a new antigen, which has been shown previously by others^31,39,40^, reflects expansion of memory B cells, leading to the production of cross neutralizing antibodies^52-54^. In addition, the XBB.1.5 booster increased neutralization titers against the advanced JN.1 variant, though titers were 5-fold lower than against XBB.1.5. This result suggests that the boosted antibodies had lower affinity or titers to cross-neutralizing determinants in JN.1. These findings are consistent with previous reports showing that JN.1 substantially evades post-vaccination sera^33,55,56^ and that the XBB.1.5 booster can provide moderate protection against JN.1^57-59^. Together, the neutralization titer data and extensive cartography analyses suggest that JN.1 may be sufficiently different from XBB.1.5 that antibody coverage could be improved with a matched vaccine antigen.

Limitations of this study include the variable time points in which convalescent sera was obtained post-COVID-19 diagnosis (between 2-59 days). The lack of human primary infection sera for Omicron variants did not allow us to confirm whether human and hamster have similar responses to these variants. In addition, cone landscapes may not be an ideal tool to predict antibody titers when newer variants such as XBB are added to the antigenic map, and we do not offer a better model. However, we bring this issue to light for further consideration of how antibody landscapes may be used to predict titers for emerging SARS-CoV-2 variants. A key strength of this study is the enhanced resolution of SARS-CoV-2 antigenic maps by including additional primary hamster sera for BA.1, BA.4/BA.5, XBB-lineage, and JN.1 variants together with a cartography approach that focuses on one antigenic cluster.

## Methods

### Cells and Viruses

The Vero E6 cell line (Cat # CRL-1586) was purchased from American Type Culture Collection (ATCC) and cultured in Dulbecco’s minimal essential medium (MEM) supplemented with 10% fetal bovine serum (FBS) (Invitrogen) and 1% penicillin/streptomycin and L-glutamine. The H1299-hACE2 cell line stably expressing human ACE2 was previously generated by lentiviral transduction of the NCI-1299 human lung carcinoma cell line (ATCC CRL-5803) with pLVX-hACE2 and selected with 1 mg/mL puromycin^60^. H1299-hACE2 cells were maintained in DMEM supplemented with 5% penicillin and streptomycin, and 10% FBS at 37°C with 5% CO2. 293T/17 cells were obtained from ATCC (#CRL-11268). 293T-ACE2-TMPRSS2 cells are available from BEI resources (#NR-55293).

SARS-CoV-2 isolates (Table S4) were obtained from BEI Resources. All virus stocks, except hCoV-19/USA-WA1/2020 (WT) and hCoV-19/USA/MD-HP24556/2022 (BA.2), were directly used for hamster infections. Due to low concentrations, the hCoV-19/USA-WA1/2020 and hCoV-19/USA/MD-HP24556/2022 were further passed on Vero E6 cells after acquiring. Passaged viruses were deep sequenced to confirm 100% match with the original sequence^61^.

### Hamster Infections

Syrian hamsters (*Mesocricetus auratus*) were acquired from Envigo (Indianapolis, IN). Animal experiments were performed in biosafety level 4 or 3 facilities accredited by the Association for Assessment and Accreditation of Laboratory Animal Care International. Studies were conducted under approved protocols (NIAID Division of Clinical Research (DCR) IRF-065E or FDA ASP 2020-06) that adhered to principles stated in the Guide for the Care and Use of Laboratory Animals, National Research Council, 2011. Primary infection sera generated at the Integrated Research Facility (NIH) used ∼6-week-old hamsters that were inoculated intranasally with 10^5^ plaque forming units (PFU) of SARS-CoV-2 in 100 μl split equally between nostrils, as previously described^62^. Primary infection sera generated at the FDA used ∼5-6 months old male hamsters inoculated with 10^4^ PFU of SARS-CoV-2 in 100 µl split equally between nostrils, as described previously^63,64^. All hamsters were subsequently monitored daily for clinical signs and weight loss. Sera were collected 7-10 days after exposures.

### Ethics statement

The PASS and EPICC clinical studies were approved by Uniformed Services University Institutional Review Board. The studies were conducted in accordance with the local legislation and institutional requirements. The participants provided informed consent to participate in this study.

### Study Cohorts

Detailed information for the serum samples is provided in Table S2. Post-vaccination sera representing different antigenic exposures were left over from our previous studies^26,29^ and came from persons enrolled in the Prospective Assessment of SARS-CoV-2 Seroconversion (PASS) study^65^. Sera from individuals who received 3 ancestral mRNA COVID-19 vaccines (V3, n=39) were collected at a median of 43 days after final vaccination dose. Sera from individuals who received 4 ancestral mRNA COVID-19 vaccines (V4, n=16) were collected at a median of 33 days after final vaccination dose. Sera from individuals who received three doses of ancestral mRNA COVID-19 vaccines and one bivalent (ancestral and BA.4/BA.5) mRNA COVID-19 vaccine dose (V3+Bi, n=19) were collected at a median of 29 days after final vaccination dose. Sera from individuals who received three ancestral mRNA COVID-19 vaccine doses before experiencing PVI during the BA.1 wave (V3+PVI, n=25) were collected at a median of 56 days after infection. Sera from persons who received four doses of ancestral mRNA COVID-19 vaccine before one bivalent (ancestral and BA.4/BA.5) mRNA COVID-19 vaccine dose (V4+Bi, n=8) were collected at a median of 37 days after final vaccination dose. Sera from persons who received three or four doses of ancestral mRNA COVID-19 vaccine, with or without a PVI before or after one bivalent (ancestral and BA.4/BA.5) mRNA COVID-19 vaccine followed by one XBB.1.5 mRNA COVID-19 vaccine (XBB.1.5 booster group*, n=16) were collected at a median of 21.5 days after the final vaccination. No subjects in V3, V4, V3+Bi, V4+Bi groups had a clinically apparent, PCR-confirmed SARS-CoV-2 infection during follow-up before serum collection. No subjects in the XBB.1.5 booster group* had a clinically apparent, PCR-confirmed SARS-CoV-2 infection after the XBB.1.5 vaccine. Genotypes for confirmed PVI are provided in Table S2.

Details of the human primary infection sera (Tables S5-S8) and neutralization titers were reported previously^26^. Most samples came from participants in the Epidemiology, Immunology, and Clinical Characteristics of Emerging Infectious Diseases with Pandemic Potential (EPICC) study^66^. Six sera came from adults following COVID-19 infection in the Republic of South Africa during a period when the Beta variant was dominant, but those infections were not genotyped, as reported previously^26^. Additionally, convalescent sera from donors with genotyped SARS-CoV-2 infections that were not represented in the EPICC study were purchased from Boca Biolistics (Pompano Beach, FL), as reported previously^26^.

### Pseudovirus Neutralization Assay

Lentiviral pseudoviruses with desired SARS-CoV-2 spike proteins (Table S4) were generated as previously described^67^. Briefly, pseudoviruses were produced in 293T cells by co-transfection of 5 μg of pCMVΔR8.2, 5 μg of pHR’CMVLuc, and 0.5 μg of pVRC8400 encoding a codon-optimized spike gene. Pseudovirus supernatants were harvested approximately 48 hours post transfection, filtered through a 0.45-μm low protein binding filter, and stored at −80°C. Pseudovirus neutralization assays were performed using 293T-ACE2-TMPRSS2 cells (BEI, NR-55293). Pseudoviruses with titers of approximately 10^6^ relative luminescence units per milliliter (RLU/mL) of luciferase activity were incubated with serially diluted sera for 2 hours at 37°C prior to inoculation onto 96-well plates that were pre-seeded one day earlier with 3.0 × 10^4^ cells per well. Pseudovirus infectivity was determined 48 hours post inoculation for luciferase activity by luciferase assay reagent (Promega) according to the manufacturer’s instructions. The inverse of the sera dilutions causing a 50% reduction of RLU compared to control was reported as the neutralization titer (ID_50_). Titers were calculated using a nonlinear regression curve fit (GraphPad Prism Software, La Jolla, California). The mean titer from at least 2 independent experiments each with intra-assay duplicates was reported as the final titer.

### Phylogenetic tree of variants in primary human infections, human post-vaccine infections, hamster inoculations and pseudoviruses

Whole genome sequences were assigned Pango lineages and amino acid substitutions and deletions were determined using Wuhan-Hu-1/2019 (NCBI accession: MN908947) as a reference. Whole genome nucleotide sequences were aligned with NextClade^68^. IQ-Tree (version 2.0.3)^69^ was used to automatically select the best-fit model and generate a consensus maximum likelihood phylogenetic tree using 1,000 bootstrap replicates. The phylogenetic tree and spike changes visualization were generated using the gheatmap function within ggtree^70^. The Genbank accession details are summarized in Table S4.

### Antigenic cartography and antibody landscapes

We used the Racmacs package (https://acorg.github.io/Racmacs/) for antigenic cartography analyses^28^ and followed the general approach of Roessler and Netzel et al. to generate antibody landscapes as described in our previous report^26^.

### Statistical Analysis

Mann–Whitney test was used for 2-group comparisons. Dunn’s multiple comparisons test for multiple groups, Pearson correlation and geometric mean titers (GMTs) with 95% confidence intervals (CIs) were determined using GraphPad Prism software. *P* values < 0.05 were considered statistically significant. All neutralization titers were log2 transformed for analyses.

### Reporting summary

Further information on research design is available in the Nature Portfolio Reporting Summary linked to this article.

## Data availability

All neutralization data are provided in the Supplemental data file. SARS-CoV-2 sequences obtained from study participants, including their accession numbers, is provided in the Supplemental tables. Data for the PASS and EPICC studies are available from the Infectious Disease Clinical Research Program (IDCRP), headquartered at the Uniformed Services University of the Health Sciences (USU), Department of Preventive Medicine and Biostatistics. Review by the USU Institutional Review Board is required for use of the data collected under these clinical protocols. Furthermore, the clinical data set includes Military Health System data collected under a Data Use Agreement that requires accounting for uses of the data. Clinical data requests may be sent to: Address: 6270A Rockledge Drive, Suite 250, Bethesda, MD 20817.

## Code availability

This study used publicly available packages and code in R, which are described in the Methods. The original R code for making the figures will be deposited at Zenodo at the time of publication.

## Material availability

De-identified serum samples from the PASS and EPICC studies are subject to a materials transfer agreement (MTA) and sample availability. Clinical sample requests may be sent to: Address: 6270A Rockledge Drive, Suite 250, Bethesda, MD 20817. Plasmid requests should be addressed to Carol Weiss and are subject to an MTA.

## Acknowledgements

We thank the White Oak FDA Animal Program and IRF-Frederick Comparative Medicine staffs for their assistance in this study.

The authors wish to also acknowledge all who have contributed to the EPICC COVID-19 study:

*Brooke Army Medical Center, Fort Sam Houston, TX:* J. Cowden; M. Darling; S. DeLeon; D. Lindholm; A. Markelz; K. Mende; S. Merritt; T. Merritt; N. Turner T. Wellington

*Carl R. Darnall Army Medical Center, Fort Hood, TX:* S. Bazan; P.K Love

*Alexander T. Augusta Military Medical Center, Fort Belvoir, VA:* N. Dimascio-Johnson; N. Elnahas; E. Ewers; K. Gallagher; C. Glinn; U. Jarral; D. Jennings; D. Larson; K. Reterstoff; A. Rutt; A. Silva; C. West; H. Al-Eid

*Henry M. Jackson Foundation, Inc.,* Bethesda*, MD:* P. Blair; J. Chenoweth; D. Clark

*Madigan Army Medical Center, Joint Base Lewis McChord, WA:* J. Bowman; S. Chambers; C. Colombo; R. Colombo; C. Conlon; K. Everson; P. Faestel; T. Ferguson; L. Gordon; S. Grogan; S. Lis; M. Martin; C. Mount; D. Musfeldt; D. Odineal; M.

Perreault; W. Robb-McGrath; R. Sainato; C. Schofield; C. Skinner; M. Stein; M. Switzer; M. Timlin; S. Wood

*Naval Medical Center* Portsmouth, Portsmouth*, VA:* S. Banks; R. Carpenter; L. Kim; K. Kronmann; T. Lalani; T. Lee; A. Smith; R. Smith; R. Tant; T. Warkentien

*Naval Medical Center San Diego, San Diego, CA:* C. Berjohn; S. Cammarata; N. Kirkland; D. Libraty; R. Maves; G. Utz

*Tripler Army Medical Center, Honolulu, HI:* C. Bradley; S. Chi; R. Flanagan; A. Fuentes; M. Jones; N. Leslie; C. Lucas; C. Madar; K. Miyasato; C. Uyehara

*Uniformed Services University of the Health Sciences,* Bethesda*, MD:* H. Adams; B. Agan; L. Andronescu; A. Austin; B. Barton, D. Becher, C. Broder; T. Burgess; C. Byrne; K Chung; J. Davies; C. English; N. Epsi; C. Fox; M. Fritschlanski; A. Hadley; P. Hickey; E. Laing; C. Lanteri; J. Livezey; A. Malloy; A. Michel, R. Mohammed; C. Morales; P. Nwachukwu; C. Olsen; E. Parmelee; S. Pollett; S. Richard; J. Rothenberg, J. Rozman; J. Rusiecki; D. Saunders; E. Samuels; M. Sanchez; A. Scher; M. Simons; A. Snow; K. Telu; D. Tribble; M. Tso; L. Ulomi; M. Wayman, N. Hockenbury

*United States Air Force School of Aerospace Medicine, Dayton, OH:* T. Chao; R. Chapleau; M. Christian; A. Fries; C. Harrington; V. Hogan; S. Huntsberger; K. Lanter; E. Macias; J. Meyer; S. Purves; K. Reynolds; J. Rodriguez; C. Starr

*United States Coast Guard, Washington, DC:* J. Iskander; I. Kamara

*Womack Army Medical Center, Fort Liberty, NC:* B. Barton; D. Hostler; J. Hostler; K. Lago; C. Maldonado; J. Mehrer

*William Beaumont Army Medical Center, El Paso, TX:* T. Hunter; J. Mejia; R. Mody; J. Montes; R. Resendez; P. Sandoval

*Walter Reed National Military Medical Center,* Bethesda*, MD:* I. Barahona; A. Baya; A. Ganesan; N. Huprikar; B. Johnson

*Walter Reed Army Institute of Research, Silver Spring, MD:* S. Peel

The authors wish to acknowledge the following individuals for their contributions to the PASS (IDCRP-126) COVID-19 study:

*Uniformed Services University (USU) Department of Microbiology and Immunology (MIC):* Eric Laing, Christopher Broder, Hannah Haines, Alyssa Lindrose, Matthew Moser, Belinda Jackson-Thompson

*USU Translational Medicine Unit:* David Saunders, Milissa Jones, Roshila Mohammed,, Priscilla Kobi, Dutchabong Shaw, Heidi Adams, and Raquel Martinez.

*USU Infectious Diseases Clinical Research Program (IDCRP):* Robert O’Connell, Timothy Burgess, Mark Simons, David Tribble, Julian Davies, Luca Illinik, Mimi Sanchez, Jennifer Rothenberg, Orlando Ortega, Edward Parmelee, Lakeesha Kosh, Bolatito Balogun

*Naval Medical Research Center - Clinical Trials Center:* Monique Hollis-Perry, Santina E. Maiolatesi, Christopher A. Duplessis, Greg Wang, Kathleen F. Ramsey, Anatalio E. Reyes, Yolanda Alcorta, and Mimi A. Wong.

*Naval Medical Research Laboratory - BDRD:* Kimberly Bishop-Lilly.

## Author contributions

Conceptualization: C.D.W., W.W.

Investigation: W.W., S.Paz, R.V., and S.L. performed all the neutralization assays; C.B.S, P.S.,T.T.W., Y.C., and M.R.H performed all the hamster infection experiments; E.G processed and managed PASS serum collections, S.D.P, B.K.A, E.M, T.H.B.

Analysis: W.W., C.D.W, G.B., R.S., L.C.K., A.F., C.K.F., M.A.C., S.A.R., N.J.E., K.B-L.

Visualization: W.W., G.B., L.C.K, C.D.W., C.K.F, M.A.C, E.G.

Writing original draft: W.W, G.B, L.C.K., and C.D.W.

Manuscript review and editing: all authors.

## Supplemental information

**Figure S1. Maximum likelihood phylogenetic tree and spike amino acid sequences used for antigenic cartography**. Taxa are color-coded for pseudovirus (PSV) (yellow) and virus in human infection sera from EPICC (Conv) (blue), PASS (PAS) (lavender), and hamster virus infection (HIF) (red). Taxa that were used for both human infection and hamster virus infection are orange. The matrix then includes spike changes relative to Wuhan-1/2019 (NCBI accession: MN908947). Deletions of multiple amino acids have been collapsed (e.g., “DEL143.145”).

**Figure S2. Neutralization titers and specificity against SARS-CoV-2 variants by human and hamster primary infection sera.** Human and hamster primary infection sera neutralization titers represented as 50% inhibitory dilutions (ID_50_) against pseudoviruses bearing spike proteins from the indicated variants are plotted. Lines indicate the results from individual samples that are connected among titers against different variants. The serum samples are the same as those in Figure 1. All data of human primary infection sera were reported previously^26^.

**Figure S3. Evaluation of goodness of fit and dimensionality for antigenic maps made with human primary infection sera (left column), hamster primary infection sera (middle column), and merged human-hamster primary infection sera (right column) related to Figure 3**. Viruses are represented by colored circles, and sera are represented by squares colored by their infecting variant. (**A**) Antigenic map with error lines. Each pair of error lines indicates the difference between the map distance and measured titer. The distance between corresponding red lines indicates when map distances are less than measured titers and blue lines indicate when map distances are greater than map titers. (**B**) Map fit of the data. Map fit was calculated by comparing the map distance to the measured titer. (**C**) Results of dimensionality testing. Cross-validation (excluding 10% of titers as a test set in 100 independent repeats) was used to determine the optimal number of dimensions. Lower root mean squared error (RMSE) for both detectable titers (above the assay limit of detection) and undetectable (below the assay limit of detection) indicate the optimal number of dimensions for fitting the antigenic map. (**D**) Maps plotted in three dimensions.

**Figure S4. Evaluation of robustness in positioning for viruses and sera on antigenic maps made with human primary infection sera (left column), hamster primary infection sera (middle column), and merged human-hamster primary infection sera (right column) related to** Figure 3. Viruses are represented as colored shapes, and sera are represented as shapes with outlines colored by corresponding infecting variant. Colors correspond to the variants labeled in Figure S3. Each grid-square represents a two-fold dilution in neutralization and is represented the same as in previous maps. (**A**) Triangulation/coordination confidence intervals for geometric uncertainty. Antigenic maps in Figure 3 show the optimal location that a virus or sera should be on a map. The coordination confidence intervals here show the broader region where the virus or sera could be placed on the map without increasing total stress of the map above one antigenic unit. (**B**) Noisy bootstrap. Random titer error (accounts for general neutralization assay variation) and antigenic noise (variation between viruses themselves) were added to the data to see how additional experimental error modifies positioning antigens on the map. (**C**) Resampled bootstrap. Some entries were resampled multiple times or completely left out to evaluate how the map would differ with alternative sampling from the underlying database.

**Figure S5. Comparison of virus positions using different cartography approaches. (A)** Procrustes analyses comparing merged human-hamster map in three dimensions to merged human-hamster map in two dimensions. Black lines point from plotted 3D map to 2D map. Each grid-square side corresponds to a 2-fold dilution in the pseudovirus neutralization assay. Antigenic distance is measured in any direction on the grid. Antigens are shown as circles and labeled. Sera are shown as squares and are colored by infecting variant. Colors correspond to the variants labeled in Figure S3. **(B)** Procrustes analyses comparing full primary infection hamster sera to pre-Omicron cluster hamster sub-map. Black arrows point from full hamster map to pre-Omicron sub map.

**Figure S6. Neutralization of the variants by sera from the different multiple antigen exposure groups.** (**A**) Amino acid mutations and deletions (Del) in spike proteins of D614G, BA.4/BA.5 and recently emerged BA.2.86-lineage and XBB-lineage variants are indicated in reference to the SARS-CoV-2/human/USA/USA-WA1/2020 (WA1/2020) (Genbank ON311289). Blue boxes indicate an amino acid substitution relative to WA1/2020. Amino acid substitutions, indicated by their single letter abbreviation, are listed in the blue box for variants that have different substitutions in those positions. N-terminal domain (NTD) and receptor binding domain (RBD) in S1 are marked. (**B**) Neutralizing antibody titers (represented as 50% inhibitory dilutions (ID_50_)) against the indicated variants in human serum samples after different antigen exposures were measured in lentiviral-based pseudovirus neutralization assays. Dots indicate results from individual participants, and bars indicate GMT with 95% confidence interval. Neutralization titers against D614G, BA.4/BA.5, BQ.1.1, XBB and XBB.1.5 in the sera of V3+PVI, V4, V3+Bi and V4+Bi were reported previously^29^.

**Figure S7. Neutralization of XBB.1.5 booster sera* subgroups based on post-vaccination infection (PVI) histories.** Neutralizing antibody titers (represented as 50% inhibitory dilutions, ID_50_) against the indicated variants in XBB.1.5 booster human serum samples with different PVIs were measured in lentiviral-based pseudovirus neutralization assays. Dots indicate results from individual participants, and bars indicate GMT with 95% confidence intervals.

**Figure S8. Antibody landscapes of multiple antigen exposure sera plotted above the XBB cluster map.** (**A–E**) Antibody landscapes are shown for individuals with (**A**) V3+Bi vaccination sera, **(B)** V3+PVI sera, **(C)** V4 vaccination sera, **(D)** V4+Bi vaccination sera, **(E)** XBB.1.5 booster sera*. (**F**) Each multiple antigen exposure serum group is stacked on top of one another for visualization. Residuals between measured and predicted titers are represented by a dotted line above or below the landscape corresponding to each antigen.

**Table S1. Antigenic distances from variants to D614G.** 95% confidence intervals were estimated by building maps with only a subset of the data and measuring the distribution of distances between each D614G and each virus across these maps.

**Table S2. Demographic information of multiple antigen exposure sera.**

**Table S3. Residual titers for all antigens in each multiple antigen exposure serum group.**

**Table S4. Viruses used for hamster infections and spikes used for pseudoviruses.**

**Table S5. Demographic information of primary infection serum samples from the EPICC study.**

**Table S6. Demographic information of primary infection serum samples commercially obtained.**

**Table S7. SARS-CoV-2 variant spikes of human primary infection serum samples from the EPICC study.**

**Table S8. SARS-CoV-2 variant spikes of human primary infection serum samples commercially obtained.**

