## Supplemental figures for "Antigenic cartography using hamster sera identifies SARS-CoV-2 JN.1 evasion seen in human XBB.1.5 booster sera"

### Supplemental Figure S1

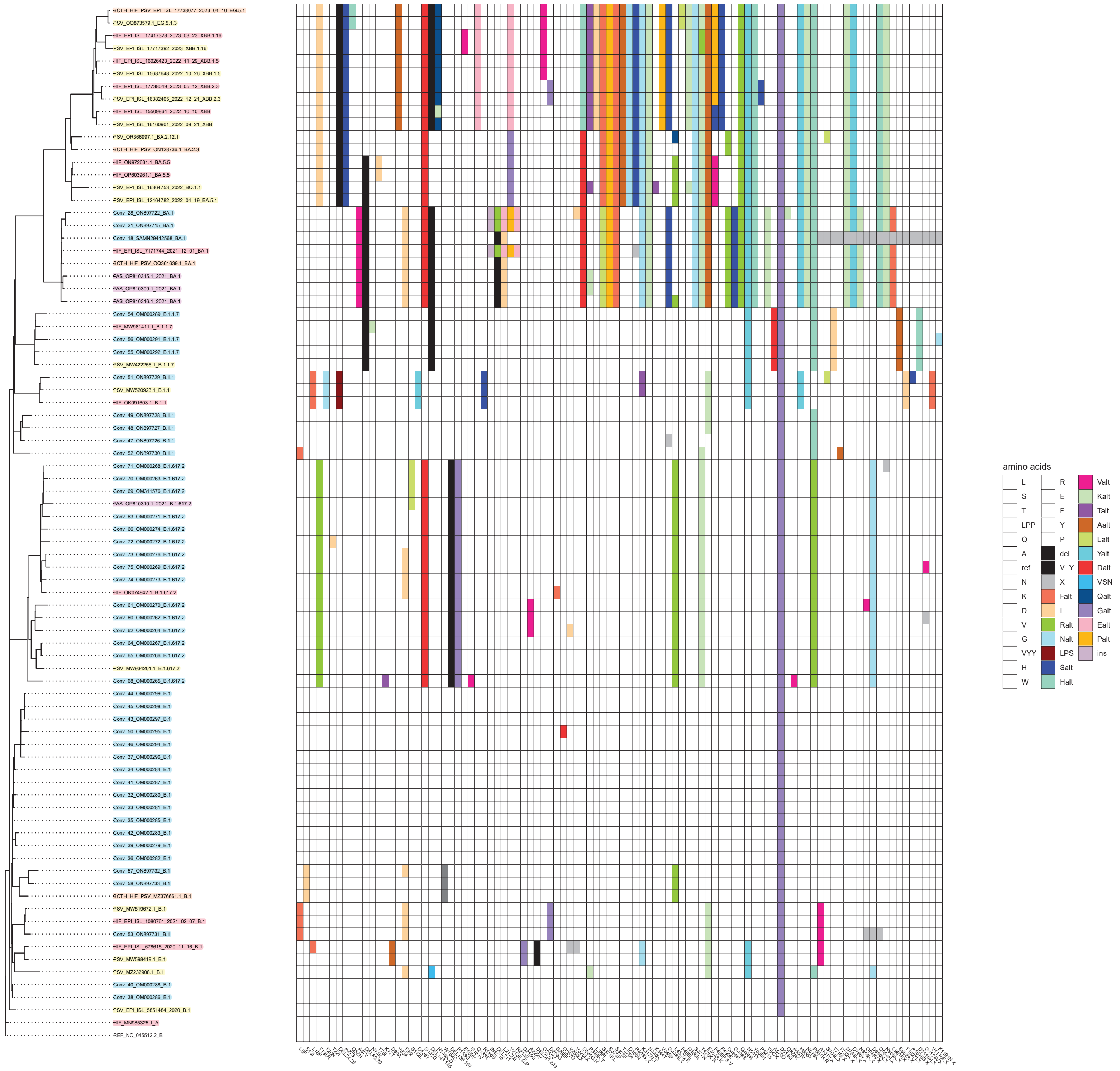


**Figure S1. Maximum likelihood phylogenetic tree and spike amino acid sequences used for antigenic cartography.** Taxa are color-coded for pseudovirus (PSV) (yellow) and virus in human infection sera from EPICC (Conv) (blue), PASS (PAS) (lavender), and hamster virus infection (HIF) (red). Taxa that were used for both human infection and hamster virus infection are orange. The matrix then includes spike changes relative to Wuhan-1/2019 (NCBI accession: MN908947). Deletions of multiple amino acids have been collapsed (e.g., "DEL143.145").

### Supplemental Figure S2

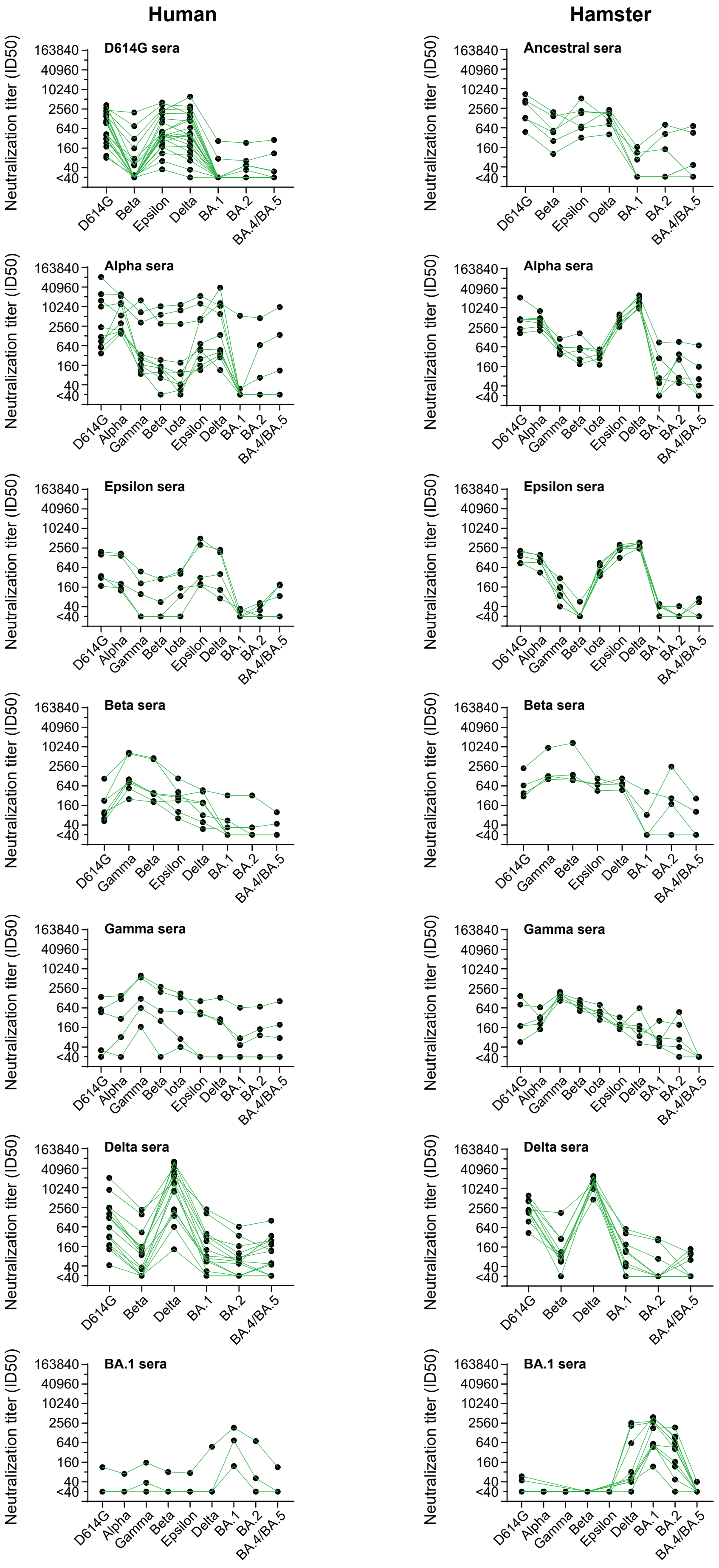

**Figure S2. Neutralization titers and specificity against SARS-CoV-2 variants by human and hamster primary infection sera.** Human and hamster primary infection sera neutralization titers represented as 50% inhibitory dilutions (ID50) against pseudoviruses bearing spike proteins from the indicated variants are plotted. Lines indicate the results from individual samples that are connected among titers against different variants. The serum samples are the same as those in Figure 1. All data of human primary infection sera were reported previously (Wang, W. *et al*. Cell Host Microbe 30, 1745-1758 e1747 (2022)).

### Supplemental Figure S3

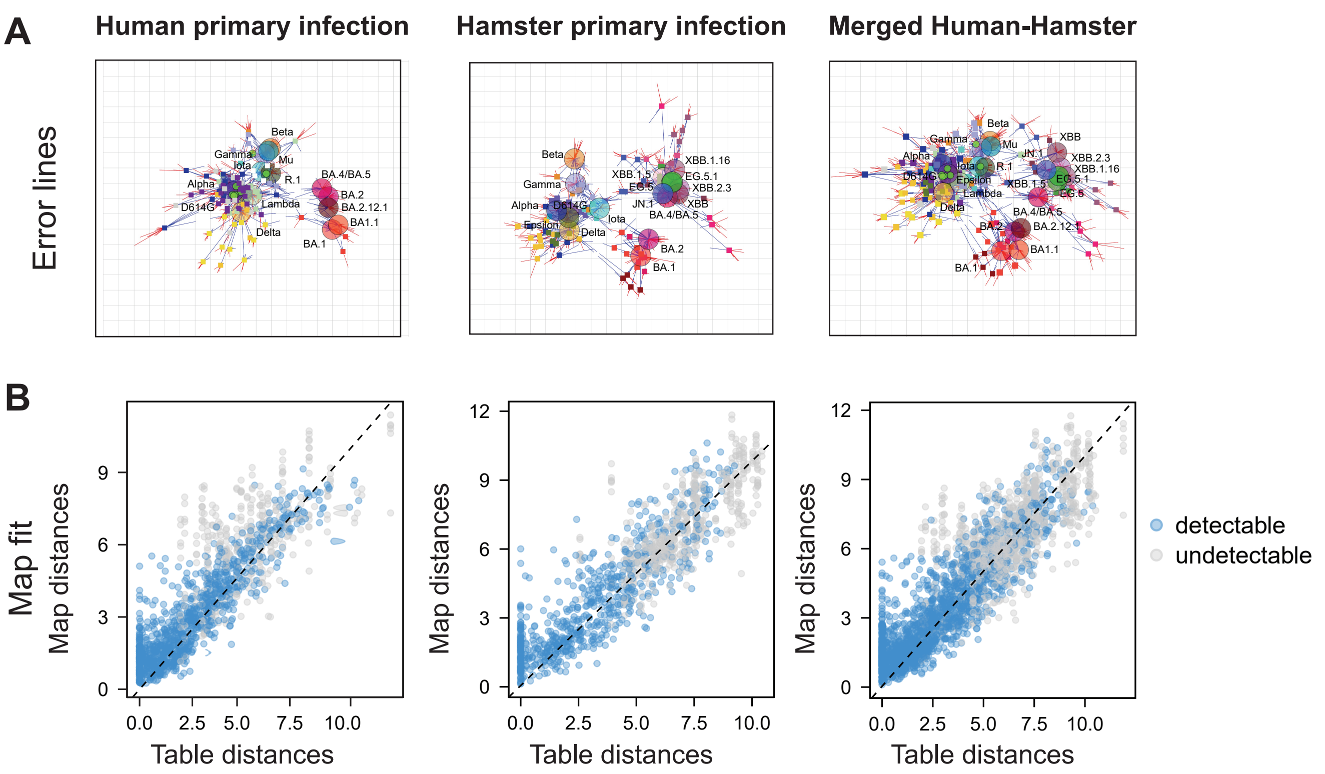


C

Dimensionality

| Dimension | Detectable Mean RMSE | Detectable Var. RMSE | Undetectable Mean RMSE | Undetectable Var. RMSE |
|---|---|---|---|---|
| 1 | 1.66 | 0.09 | 2.08 | 0.11 |
| 2 | 1.41 | 0.06 | 1.95 | 0.07 |
| 3 | 1.41 | 0.08 | 1.78 | 0.08 |
| 4 | 1.43 | 0.09 | 1.74 | 0.07 |
| 5 | 1.43 | 0.10 | 1.75 | 0.14 |

| Dimension | Detectable Mean RMSE | Detectable Var. RMSE | Undetectable Mean RMSE | Undetectable Var. RMSE |
|---|---|---|---|---|
| 1 | 2.41 | 0.13 | 2.21 | 0.11 |
| 2 | 2.10 | 0.09 | 1.48 | 0.09 |
| 3 | 2.06 | 0.12 | 1.26 | 0.06 |
| 4 | 2.08 | 0.12 | 1.20 | 0.06 |
| 5 | 2.08 | 0.14 | 1.16 | 0.07 |

| Dimension | Detectable Mean RMSE | Detectable Var. RMSE | Undetectable Mean RMSE | Undetectable Var. RMSE |
|---|---|---|---|---|
| 1 | 2.30 | 0.10 | 2.20 | 0.10 |
| 2 | 1.77 | 0.11 | 1.67 | 0.06 |
| 3 | 1.74 | 0.10 | 1.48 | 0.04 |
| 4 | 1.76 | 0.11 | 1.41 | 0.03 |
| 5 | 1.77 | 0.14 | 1.41 | 0.06 |

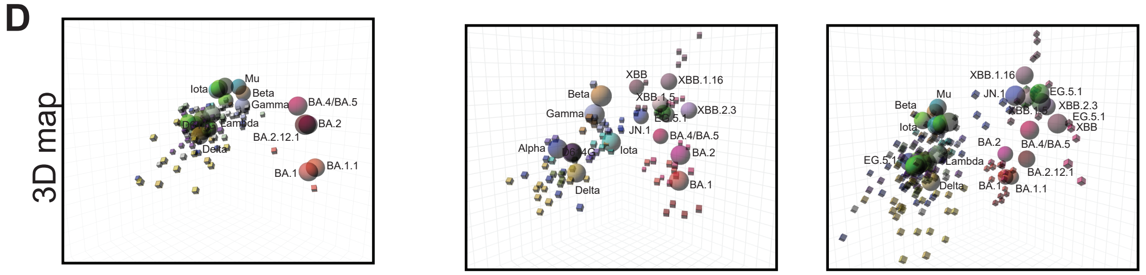


**Figure S3. Evaluation of goodness of fit and dimensionality for antigenic maps made with human primary infection sera (left column), hamster primary infection sera (middle column), and merged human-hamster primary infection sera (right column) related to Figure 3.** Viruses are represented by colored circles, and sera are represented by squares colored by their infecting variant. (**A**) Antigenic map with error lines. Each pair of error lines indicates the difference between the map distance and measured titer. The distance between corresponding red lines indicate when map distances are less than measured titers and blue lines indicate when map distances are greater than map titers. (**B**) Map fit of the data. Map fit was calculated by comparing the map distance to the measured titer. (**C**) Results of dimensionality testing. Cross-validation (excluding 10% of titers as a test set in 100 independent repeats) was used to determine the optimal number of dimensions. Lower root mean squared error (RMSE) for both detectable titers (above the assay limit of detection) and undetectable (below the assay limit of detection) indicate the optimal number of dimensions for fitting the antigenic map. (**D**) Maps plotted in three dimensions.

### Supplemental Figure S4

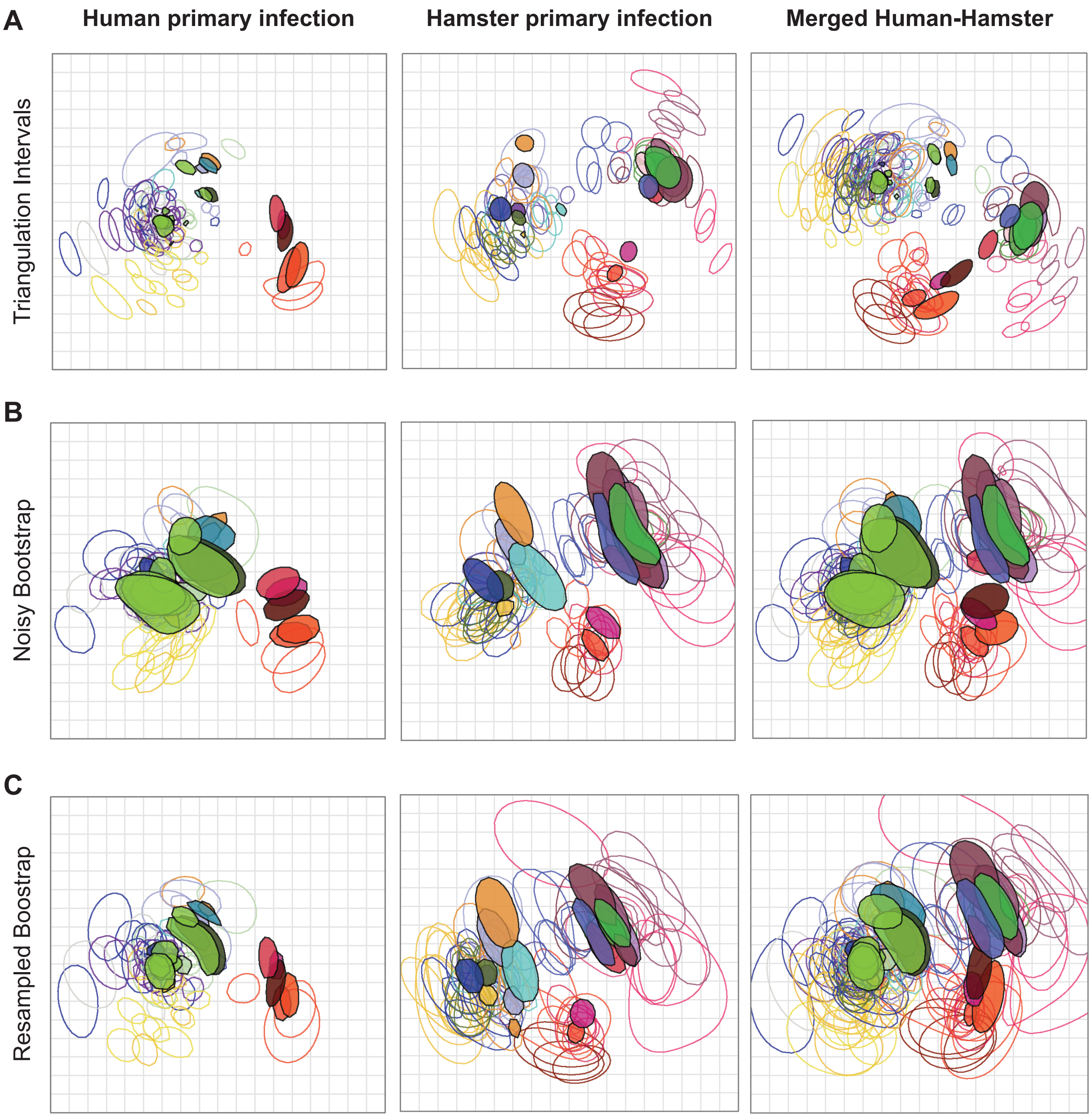


**Figure S4. Evaluation of robustness in positioning for viruses and sera on antigenic maps made with human primary infection sera (left column), hamster primary infection sera (middle column), and merged human-hamster primary infection sera (right column) related to Figure 3.** Viruses are represented as colored shapes, and sera are represented as shapes with outlines colored by corresponding infecting variant. Colors correspond to the variants labeled in Figure S3. Each grid-square represents a two-fold dilution in neutralization and is represented the same as in previous maps. (**A**) Triangulation/coordination confidence intervals for geometric uncertainty. Antigenic maps in Figure 3 show the optimal location that a virus or sera should be on a map. The coordination confidence intervals here show the broader region where the virus or sera could be placed on the map without increasing total stress of the map above one antigenic unit. (**B**) Noisy bootstrap. Random titer error (accounts for general neutralization assay variation) and antigenic noise (variation between viruses themselves) were added to the data to see how additional experimental error modifies positioning antigens on the map. (**C**) Resampled bootstrap. Some entries were resampled multiple times or completely left out to evaluate how the map would differ with alternative sampling from the underlying database.

Supplemental Figure S5

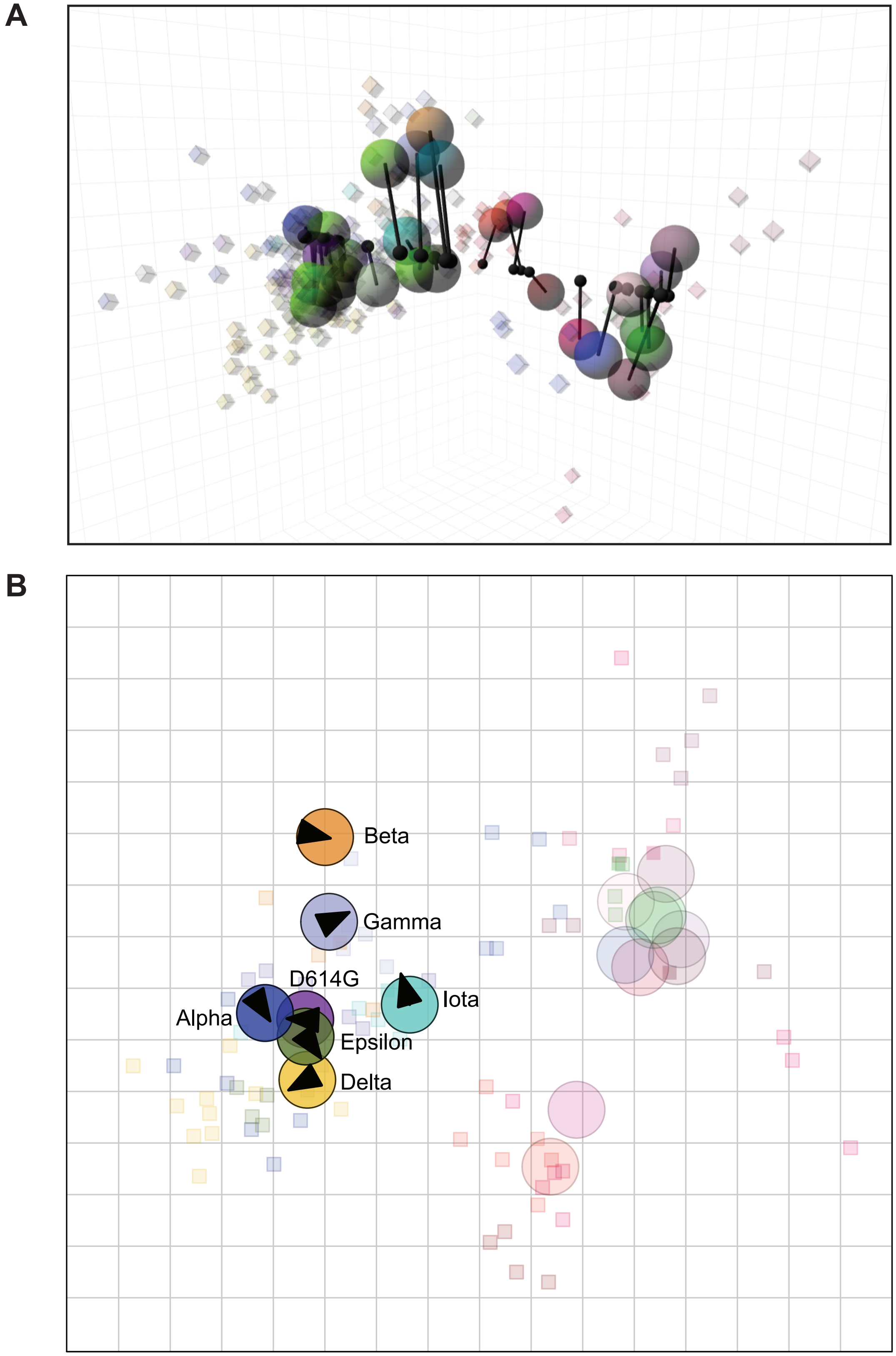


**Figure S5. Comparison of virus positions using different cartography approaches.** (**A**) Procrustes analyses comparing merged human-hamster map in three dimensions to merged human-hamster map in two dimensions. Black lines point from plotted 3D map to 2D map. Each grid-square side corresponds to a 2-fold dilution in the pseudovirus neutralization assay. Antigenic distance is measured in any direction on the grid. Antigens are shown as circles and labeled. Sera are shown as squares and are colored by infecting variant. Colors correspond to the variants labeled in Figure S3. (**B**) Procrustes analyses comparing full primary infection hamster sera to pre-Omicron cluster hamster sub-map. Black arrows point from full hamster map to pre-Omicron sub map.

### Supplemental Figure S6

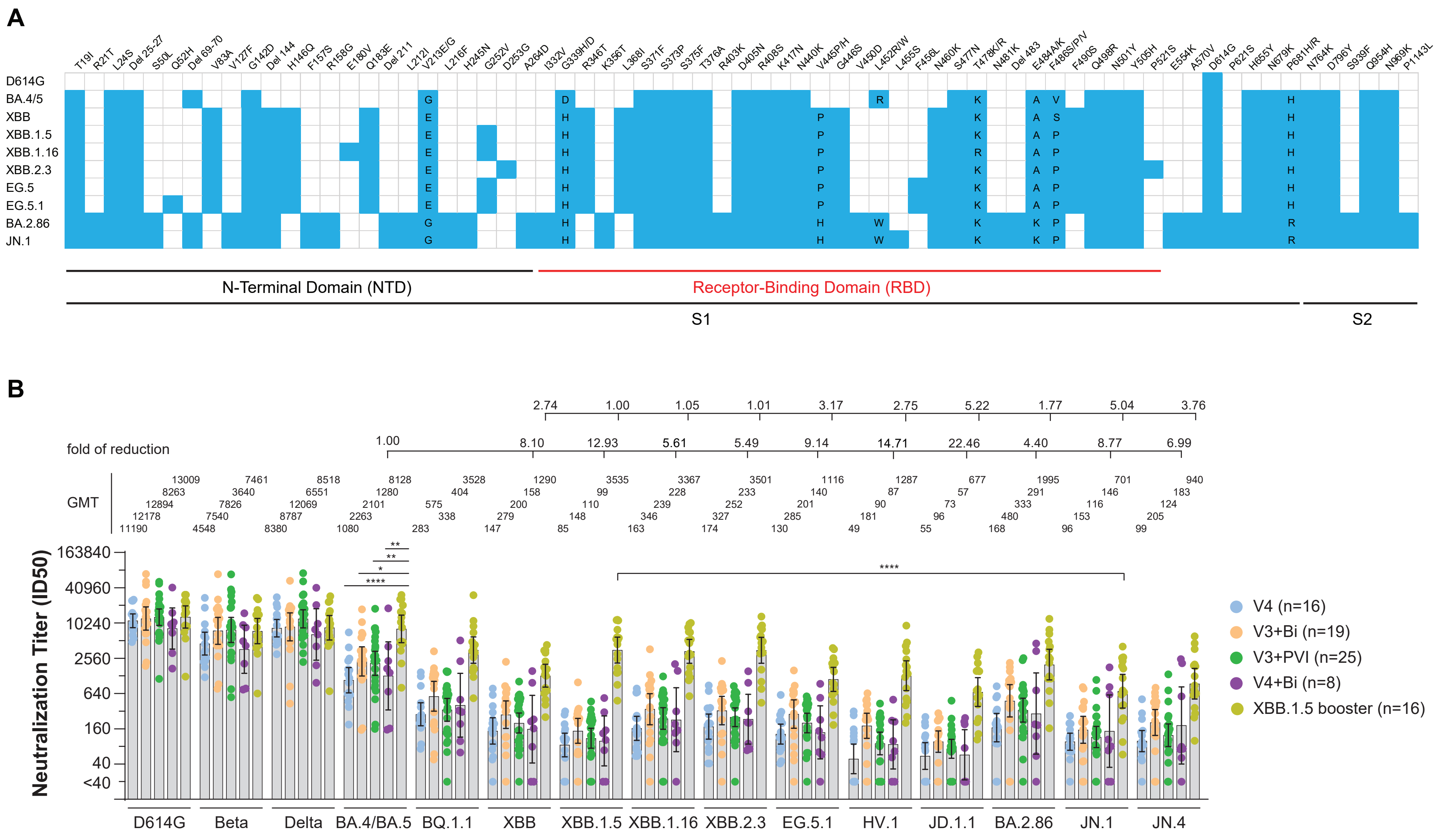

**Figure S6. Neutralization of the variants by sera from the different multiple antigen exposure groups.** (**A**) Amino acid mutations and deletions (Del) in spike proteins of D614G, BA.4/BA.5 and recently emerged BA.2.86-lineage and XBB-lineage variants are indicated in reference to the SARS-CoV-2/human/USA/USA-WA1/2020 (WA1/2020) (Genbank ON311289). Blue boxes indicate an amino acid substitution relative to WA1/2020. Amino acid substitutions, indicated by their single letter abbreviation, are listed in the blue box for variants that have different substitutions in those positions. N-terminal domain (NTD) and receptor binding domain (RBD) in S1 are marked. (**B**) Neutralizing antibody titers (represented as 50% inhibitory dilutions (ID50)) against the indicated variants in human serum samples after different antigen exposures were measured in lentiviral-based pseudovirus neutralization assays. Dots indicate results from individual participants, and bars indicate GMT with 95% confidence interval. Neutralization titers against D614G, BA.4/BA.5, BQ.1.1, XBB and XBB.1.5 in the sera of V3+PVI, V4, V3+Bi and V4+Bi were reported previously (Wang, W. *et al*. J Infect Dis 228, 439-443 (2023)).

### Supplemental Figure S7

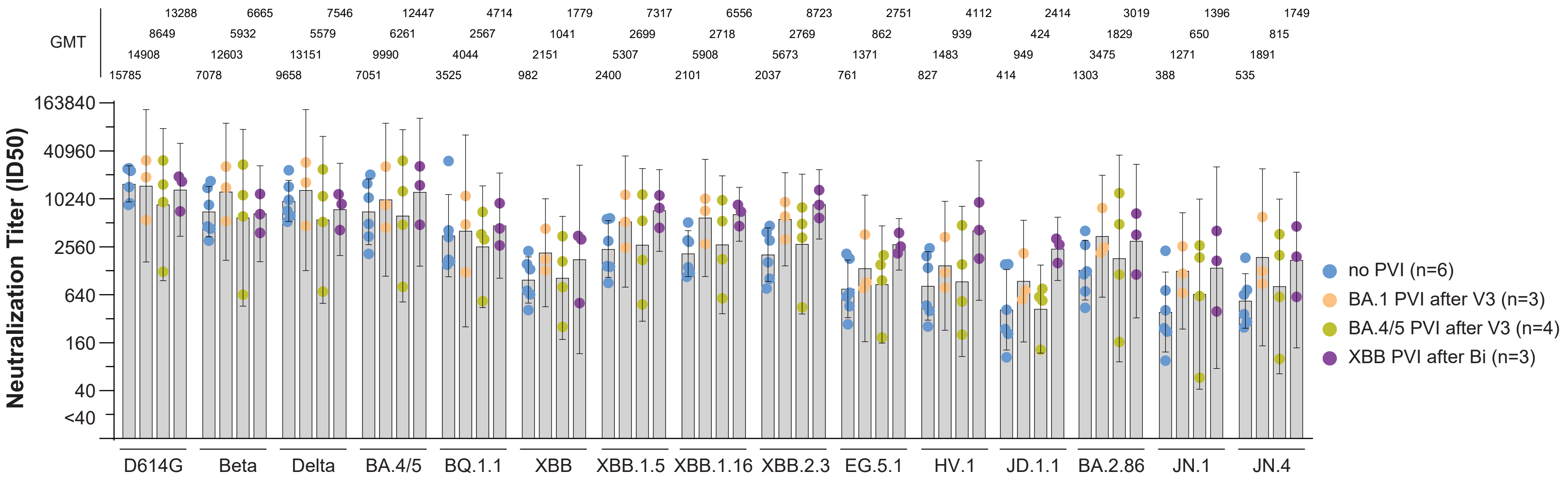


**Figure S7. Neutralization of XBB.1.5 booster sera* subgroups based on post-vaccination infection (PVI) histories.** Neutralizing antibody titers (represented as 50% inhibitory dilutions, ID50) against the indicated variants in XBB.1.5 booster human serum samples with different PVIs were measured in lentiviral-based pseudovirus neutralization assays. Dots indicate results from individual participants, and bars indicate GMT with 95% confidence intervals.

### Supplemental Figure S8

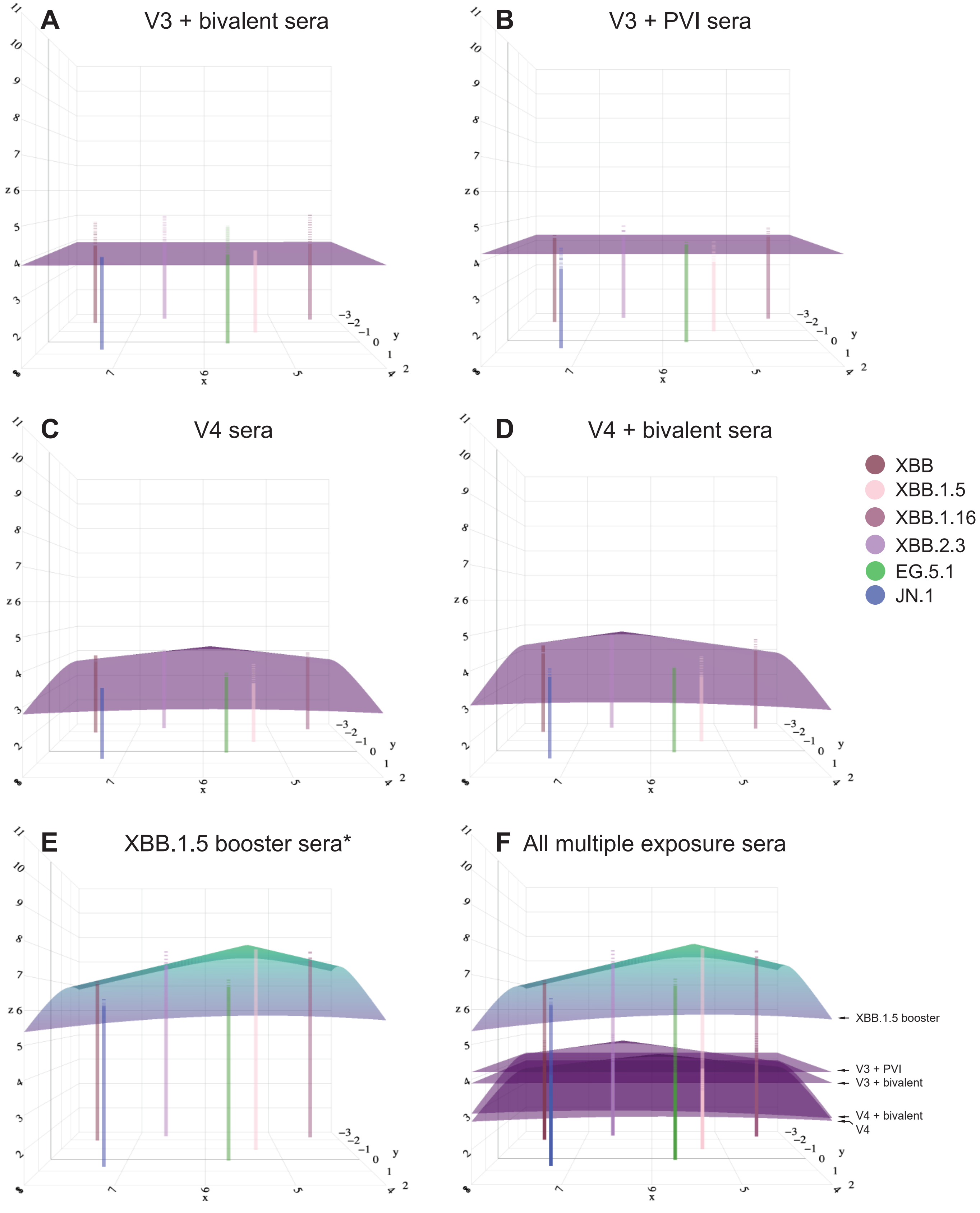


**Figure S8. Antibody landscapes of multiple antigen exposure sera plotted above the XBB cluster map.** (**A–E**) Antibody landscapes are shown for individuals with (**A**) V3+Bi vaccination sera, (**B**) V3+PVI sera, (**C**) V4 vaccination sera, (**D**) V4+Bi vaccination sera, (**E**) XBB.1.5 booster sera*. (**F**) Each multiple antigen exposure serum group is stacked on top of one another for visualization. Residuals between measured and predicted titers are represented by a dotted line above or below the landscape corresponding to each antigen.
