## Supplemental tables for "Antigenic cartography using hamster sera identifies SARS-CoV-2 JN.1 evasion seen in human XBB.1.5 booster sera"

**Table S1. Antigenic distances from variants to D614G**

| | Human map (n=84) | Hamster map (n=82) | Merged human-hamster map (n=166) |
|---|---|---|---|
| **Alpha** | 0.6 (0.4 – 0.7) | 0.9 (0.6 – 1.3) | 0.7 (0.6 – 0.9) |
| **Beta** | 3.8 (3.7 – 3.9) | 3.6 (3.3 – 3.8) | 3.6 (3.4 – 3.7) |
| **Gamma** | 3.3 (3.2 – 3.4) | 2.1 (1.8 – 2.4) | 3.1 (3.0 – 3.2) |
| **Iota** | 2.3 (2.2 – 2.5) | 2.1 (1.8 – 2.3) | 2.1 (2.0 – 2.3) |
| **Epsilon** | 0.6 (0.5 – 0.8) | 0.2 (0.0 – 0.5) | 0.3 (0.2 – 0.4) |
| **Delta** | 1.5 (1.4 -1.6) | 1.2 (1.0 – 1.4) | 1.4 (1.3 – 1.5) |
| **Lambda** | 1.3 (1.2 – 1.5) | - | 1.1 (0.9 – 1.2) |
| **Mu** | 3.6 (3.5 – 3.8) | - | 3.4 (3.3 – 3.6) |
| **BA.1** | **7.0 (6.9 – 7.2)** | **5.7 (5.5 – 6.0)** | 6.3 (6.2 – 6.5) |
| **BA.1.1** | 7.3 (7.2 – 7.5) | - | 7.0 (6.8 – 7.1) |
| **BA.2** | 6.4 (6.2 – 6.6) | 5.6 (5.3 – 5.9) | 6.0 (5.8 – 6.2) |
| **BA.2.12.1** | 6.4 (6.3 -6.6) | - | 6.1 (6.0 – 6.3) |
| **BA.4/BA.5** | **5.9 (5.8 – 6.1)** | **6.6 (6.3 – 7.0)** | 6.3 (6.1 – 6.5) |
| **XBB** | - | 7.2 (6.9 – 7.6) | 7.4 (7.1 – 7.7) |
| **XBB.1.16** | - | 7.7 (7.4 – 8.0) | 7.9 (7.7 – 8.3) |
| **XBB.1.5** | - | 6.7 (6.5 – 7.0) | 7.0 (6.9 – 7.2) |
| **XBB.2.3** | - | 7.4 (7.1 – 7.8) | 7.6 (7.3 – 8.0) |
| **EG.5** | - | 7.3 (7.0 – 7.5) | 7.5 (7.3 – 7.8) |
| **EG.5.1** | - | 7.1 (7.2 – 8.4) | 7.3 (7.1 – 7.6) |
| **JN.1** | - | 6.4 (6.1 – 6.7) | 6.6 (6.3 – 6.9) |

Antigenic distances (antigenic units) between D614G and variants from human primary infection map, hamster primary infection map and merged human-hamster map. An antigenic distance equivalent to one is represented by one grid-square on the antigenic map. 95% confidence intervals for antigenic distances were calculated.

**Table S2. Demographic information of multiple antigen exposure sera from the PASS study**

| | Overall (n=123)[a] | V3 (n=39) | V3 + Bi (n=19) | V3 + PVI (n=25) | V4 (n=16) | V4 + Bi (n=8) | XBB.1.5 booster (n=16)[c] |
|---|---|---|---|---|---|---|---|
| **Age group** | | | | | | | |
| 18-44 | 57 (46.3%) | 19 (48.7%) | 15 (78.9%) | 12 (48.0%) | 2 (12.5%) | 0 (0%) | 9 (56.3%) |
| 45-64 | 63 (51.2%) | 19 (48.7%) | 4 (21.1%) | 13 (52.0%) | 13 (81.3%) | 7 (87.5%) | 7 (43.7%) |
| 65+ | 3 (2.5%) | 1 (2.6%) | 0 (0%) | 0 (0%) | 1 (6.2%) | 1 (12.5%) | 0 (0%) |
| Median age (IQR) | 46.0 (38.0 - 54.0) | 45.0 (35.0 - 52.0) | 39.0 (34.5 - 43.0) | 45.0 (38.0 - 53.0) | 55.5 (51.0 - 58.0) | 54.5 (53.5 - 59.3) | 41.5 (37.8 - 50.5) |
| **Gender** | | | | | | | |
| Female | 82 (66.7%) | 25 (64.1%) | 11 (57.9%) | 18 (72.0%) | 10 (62.5%) | 6 (75.0%) | 12 (75.0%) |
| Male | 41 (33.3%) | 14 (35.9%) | 8 (42.1%) | 7 (28.0%) | 6 (37.5%) | 2 (25.0%) | 4 (25.0%) |
| **Race/Ethnicity** | | | | | | | |
| White | 80 (65.0%) | 25 (64.1%) | 14 (73.7%) | 17 (68.0%) | 9 (56.3%) | 6 (75.0%) | 9 (56.3%) |
| Black | 15 (12.2%) | 4 (10.3%) | 2 (10.5%) | 3 (12.0%) | 2 (12.5%) | 1 (12.5%) | 3 (18.7%) |
| Hispanic or Latino | 7 (5.7%) | 1 (2.6%) | 0 (0%) | 2 (8.0%) | 3 (18.8%) | 1 (12.5%) | 0 (0%) |
| Others | 21 (17.1%) | 9 (23.1%) | 3 (15.8%) | 3 (12.0%) | 2 (12.5%) | 0 (0%) | 4 (25.0%) |
| **Charlson Co-morbidity Index** | | | | | | | |
| 0 | 106 (86.2%) | 34 (87.9%) | 17 (89.5%) | 21 (84.0%) | 13 (81.2%) | 5 (62.5%) | 16 (100%) |
| 1 | 9 (7.3%) | 2 (5.1%) | 2 (10.5%) | 3 (12.0%) | 1 (6.2%) | 1 (12.5%) | 0 (0%) |
| 2 | 5 (4.0%) | 2 (5.1%) | 0 (0%) | 1 (4.0%) | 1 (6.2%) | 1 (12.5%) | 0 (0%) |
| 3 | 3 (2.5%) | 1 (2.6%) | 0 (0%) | 0 (0%) | 1 (6.2%) | 1 (12.5%) | 0 (0%) |
| **Primary vaccine (V1+V2)** | | | | | | | |
| Pfizer BNT162b2 | 123 (100%) | 39 (100%) | 19 (100%) | 25 (100%) | 16 (100%) | 8 (100%) | 16 (100%) |
| **First booster (V3)** | | | | | | | |
| Pfizer BNT162b2 | 120 (97.5%) | 39 (100%) | 19 (100%) | 24 (96.0%) | 15 (93.8%) | 8 (100%) | 15 (93.7%) |
| Moderna mRNA-1273 | 3 (2.5%) | 0 (0%) | 0 (0%) | 1 (4.0%) | 1 (6.2%) | 0 (0%) | 1 (6.3%) |
| **Second booster (V4)** | | | | | | | |
| Pfizer BNT162b2 | 22 (17.9%) | - | 0 (0%) | - | 13 (81.2%) | 7 (87.5%) | 2 (12.5%) |
| Pfizer bivalent (original and Omicron BA.4/BA.5) | 31 (25.2%) | - | 19 (100%) | - | 0 (0%) | 0 (0%) | 12 (75.0%) |
| Moderna mRNA-1273 | 4 (3.3%) | - | 0 (0%) | - | 3 (18.8%) | 1 (12.5%) | 0 (0%) |
| Moderna bivalent (original and Omicron BA.4/BA.5) | 1 (0.8%) | - | - | - | - | - | 1 (6.3%) |
| Pfizer monovalent XBB.1.5 | 1 (0.8%) | - | - | - | - | - | 1 (6.3%) |
| Unboosted | 64 (52.0%) | - | 0 (0%) | - | 0 (0%) | 0 (0%) | 0 (0%) |
| **Third booster (V5)** | | | | | | | |
| Pfizer bivalent (original and Omicron BA.4/BA.5) | 10 (8.1%) | - | - | - | - | 8 (100%) | 2 (12.5%) |
| Pfizer monovalent XBB.1.5 | 12 (9.8%) | - | - | - | - | - | 12 (75.0%) |
| Moderna monovalent XBB.1.5 | 1 (0.8%) | - | - | - | - | - | 1 (6.3%) |
| Unboosted | 100 (81.3%) | - | - | - | - | - | 1 (6.3%) |
| **Fourth booster (V6)** | | | | | | | |
| Pfizer monovalent XBB.1.5 | 2 (1.6%) | - | - | - | - | - | 2 (12.5%) |
| Unboosted | 121 (98.4%) | - | - | - | - | - | 14 (87.5%) |
| **Days between most recent vaccination and serum sample collection** | | | | | | | |
| Median (IQR) | 39.0 (27.0 - 61.0) | 43.0 (33.5 - 53.5) | 29.0 (27.5 - 35.0) | 169.0 (148.0 - 267.0) | 33.0 (22.5 - 53.0) | 37.0 (32.8 - 43.0) | 21.5 (16.0 - 27.3) |
| **Days between infection symptom onset and serum sample collection** | | | | | | | |
| Median (IQR) | 70.5 (38.8 - 184.0) | - | - | 56.0 (37.0 - 75.0) | - | - | 553.0 (446.5 - 672.0) |
| **Infecting genotype[b]** | | | | | | | |
| BA.1.1 | 3 (2.5%) | - | - | 2 (8.0%) | - | - | 1 (6.3%) |
| BA.1.18 | 1 (0.8%) | - | - | - | - | - | 1 (6.3%) |
| BA.1.19 | 1 (0.8%) | - | - | 1 (4.0%) | - | - | - |
| AY.25 | 1 (0.8%) | - | - | 1 (4.0%) | - | - | - |

[a]Samples collected from PASS study; 29 participants contributed serum samples collected at different timepoints to 2 or 3 groups (3 contributed to V3 and V4, 3 contributed to V3 and V3+Bi, 6 contributed to V3 and V3+PVI, 1 contributed to V3 and V4+Bi, 3 contributed to V3 and XBB.1.5 booster, 4 contributed to V4 and V4+Bi, 3 contributed to V3+Bi and XBB.1.5 booster, 2 contributed to V3+PVI and XBB.1.5 booster, 1 contributed to V3, V4 and V4+Bi, 1 contributed to V3, V4+Bi and XBB.1.5 booster, and 2 contributed to V3, V3+PVI and XBB.1.5 booster)

[b]Genotypes assigned based on Pango 4.1.2

IQR, Interquartile range; V1+V2, 2 doses ancestral mRNA vaccine; V3, 3 doses ancestral mRNA vaccine; V3+Bi, 3 doses ancestral + 1 dose bivalent mRNA vaccine; V4, 4 doses ancestral mRNA vaccine; V4+Bi, 4 doses ancestral + 1 dose bivalent mRNA vaccine

[c]XBB.1.5 booster group includes six individuals without reported PVI, three individuals with presumed BA.1 PVIs and four individuals with presumed BA.5 PVIs after three doses of the ancestral COVID-19 vaccine, and three individuals with presumed XBB PVIs after the bivalent booster.

**Table S3. Residual titers for all antigens in each multiple antigen exposure serum group**

| | **V3 + bivalent** (n = 19) | **V3 + PVI** (n = 23) | **V4** (n = 16) | **V4 + bivalent** (n = 8) | **XBB 1.5 Booster** (n = 16) |
|---|---|---|---|---|---|
| **D614G** | 0.48 | 0.26 | 0.36 | 0.36 | 0.23 |
| **Beta** | 1.49 | 1.72 | 1.41 | 1.3 | 0.23 |
| **Delta** | 0.71 | 0.93 | 0.73 | 0.67 | -0.01 |
| **BA.4/BA.5** | 2.45 | 2.65 | 2.2 | 2.26 | 1.66 |
| **XBB.1.16** | 0.32 | 0.19 | 0.18 | 0.41 | 0.67 |
| **XBB.1.5** | -1.5 | -1.6 | -1.46 | -1.4 | 0.45 |
| **XBB.2.3** | 0.25 | 0.26 | 0.26 | 0.42 | 0.74 |
| **XBB** | -0.03 | -0.15 | 0.01 | -0.21 | -0.73 |
| **EG.5.1** | -0.25 | -0.4 | -0.5 | -0.61 | 1.06 |
| **JN.1** | -1.59 | -1.7 | -1.46 | -1 | -1.95 |

Residual titers are calculated by subtracting the predicted titer (from fitting the landscape) from the measured titer and are represented as log titers.
Positive residual titers indicate that measured titer was greater than predicted titer. Negative residual titers indicate that measured titer was less than predicted titer.

**Table S4. Viruses used for hamster infections and spikes used for pseudoviruses.**

| Virus used for hamster infection | | | | Spike used in pseudovirus | |
|---|---|---|---|---|---|
| Name | Strain | Source | GenBank or GISAID # | Name | GenBank or GISAID # |
| WT | hCoV-19/USA-WA1/2020 | BEI, NR-52281 | MN985325.1 | D614G | EPI_ISL_5851484 |
| B.1.1.7 | hCoV-19/USA/CA_CDC_5574/2020 | BEI, NR-54011 | MW981411 | B.1.1.7 | MW422256 |
| P.1 | hCoV-19/Japan/TY7-503/2021 | BEI, NR-54984 | OK091603 | P.1 | MW520923 |
| B.1.427 | hCoV-19/USA/CA-CDC-48018/2020 | BEI, NR-55338 | MZ376661 | B.1.427 | MZ376661 |
| B.1.526 | hCoV-19/USA/NY-NP-DOH1/2021 | BEI, NR-55637 | EPI_ISL_1080761 | B.1.526 | MW519672 |
| B.1.617.2 | hCoV-19/USA/PHC658/2021 | BEI, NR-55612 | OR074942 | B.1.617.2 | MW934201 |
| BA.1 | hCoV-19/USA/MD-HP20874/2021 | BEI, NR-56461 | OQ361639 | BA.1 | OQ361639 |
| BA.1.1 | hCoV-19/USA/GA-EHC2811C/2021 | BEI, NR-56482 | EPI_ISL_7171744 | | |
| BA.2 | hCoV-19/USA/MD-HP24556/2022 | BEI, NR-56512 | ON128736 | BA.2 | ON128736 |
| BA.5 | hCoV-19/ USA/COR-22-063113/2022 | BEI, NR-58616 | ON972631 | BA.4/5 | EPI_ISL_12464782 |
| XBB.1.5 | hCoV-19/USA/MD-HP40900/2022 | BEI, NR-59104 | EPI_ISL_16026423 | XBB.1.5 | EPI_ISL_15687648 |
| XBB.1.16 | hCoV-19/USA/CA-Stanford-139_S23/2023 | BEI, NR-59442 | EPI_ISL_17417328 | XBB.1.16 | EPI_ISL_17717392 |
| | | | | XBB.2.3 | EPI_ISL_16382405 |
| | | | | BQ.1.1 | EPI_ISL_16364753 |
| XBB | hCoV-19/USA/CA-Stanford-109_S21/2022 | BEI, NR-58925 | EPI_ISL_15509864 | XBB | EPI_ISL_16160901 |
| B.1.351 | hCoV-19/South Africa/KRISP-K005325/2020 | BEI, NR-54009 | EPI_ISL_678615 | B.1.351 | MW598419 |
| | | | | B.1.621 | MZ232908 |
| | | | | BA.2.12.1 | OR366997 |
| | | | | EG.5 | OQ873579 |
| EG.5.1 | hCoV-19/USA/MD-HP47946/2023 | BEI, NR-59503 | EPI_ISL_17738077 | EG.5.1 | EPI_ISL_17738077 |

**Table S5. Demographic information of primary infection serum samples from the EPICC study**

| | Overall (n=45)[a] |
|---|---|
| **Age group** | |
| <18 | 5 (11.1%) |
| 18-44 | 20 (44.4%) |
| 45-64 | 15 (33.3%) |
| 65+ | 5 (11.1%) |
| **Gender** | |
| Female | 19 (42.2%) |
| Male | 26 (57.8%) |
| **Race/ Ethnicity** | |
| White | 21 (46.7%) |
| Hispanic or Latino | 14 (31.1%) |
| Black | 7 (15.6%) |
| Others | 3 (6.7%) |
| **Severity of initial infection** | |
| Hospitalized | 23 (51.1%) |
| Outpatient | 22 (48.9%) |
| **Charlson comorbidity index** | |
| 0 | 19 (42.2%) |
| 1-2 | 17 (37.8%) |
| 3-4 | 6 (13.3%) |
| >5 | 3 (6.7%) |
| **Primary vaccine** | |
| Unvaccinated | 45 (100.0%) |
| **Days between infection symptom onset and sera sample collection** | |
| Median ± SD (range) | 27 ± 9.7 (8.0 - 51.0) |
| **Infecting genotype**[b] | |
| B.1 | 10 (22.2%) |
| B.1.2 | 6 (13.3%) |
| BA.1 | 3 (6.7%) |
| B.1.1.207 | 3 (6.7%) |
| B.1.1.7 | 3 (6.7%) |
| AY.100 | 2 (4.4%) |
| AY.14 | 2 (4.4%) |
| AY.25 | 2 (4.4%) |
| B.1.617.2 | 2 (4.4%) |
| B.1.429 | 2 (4.4%) |
| AY.119 | 1 (2.2%) |
| AY.25.1 | 1 (2.2%) |
| AY.3 | 1 (2.2%) |
| AY.44 | 1 (2.2%) |
| AY.47 | 1 (2.2%) |
| AY.62 | 1 (2.2%) |
| AY.74 | 1 (2.2%) |
| B.1.1.519 | 1 (2.2%) |
| B.1.526 | 1 (2.2%) |
| P.1.10 | 1 (2.2%) |

[a] *Sample collected from EPICC study*

[b] *Genotypes assigned based on Pango 4.0.6*

**Table S6. Demographic information of primary infection serum samples commercially obtained**

| | Overall (N=31) |
|---|---|
| **Gender** | |
| Female | 10 (32.3%) |
| Male | 21 (67.7%) |
| **Infecting genotype** | |
| B.1 | 2 (6.5%) |
| B.1.1.7 | 7 (22.6%) |
| B.1.2 | 2 (6.5%) |
| B.1.234 | 1 (3.2%) |
| B.1.429 | 3 (9.7%) |
| B.1.577 | 1 (3.2%) |
| C.11 | 1 (3.2%) |
| C.37 | 10 (32.3%) |
| P.1 | 4 (12.9%) |
| **Time between infection symptom onset and sera sample collection** | |
| Median ± SD (range) | 5 ± 12.6 (2.0 - 59.0) |

**Table S7. SARS-CoV-2 variant spikes of human primary infection serum samples from the EPICC study**

| SampleID | Pangolin 4.0.6 | Spike Substitutions | Spike Deletions | Accession |
|---|---|---|---|---|
| Conv-15 | BA.1 | S:A67V,S:T95I,S:Y145D,S:L212I,S:G339D,S:S371L,S:S373P,S:S375F,S:K417N,S:N440K,S:G446S,S:S477N,S:T478K,S:E484A,S:Q493R,S:G496S,S:Q498R,S:N501Y,S:Y505H,S:T547K,S:D614G,S:H655Y,S:N679K,S:P681H | S:H69-,S:V70-,S:G142-,S:V143-,S:Y144-,S:N211- | SAMN29442568 |
| Conv-18 | BA.1 | S:A67V,S:T95I,S:Y145D,S:L212I,S:G339D,S:S371L,S:S373P,S:S375F,S:K417N,S:N440K,S:G446S,S:S477N,S:T478K,S:E484A,S:Q493R,S:G496S,S:Q498R,S:N501Y,S:Y505H,S:T547K,S:D614G,S:H655Y,S:N679K,S:P681H,S:N764K,S:D796Y,S:N856K,S:Q954H,S:N969K,S:L981F | S:H69-,S:V70-,S:G142-,S:V143-,S:Y144-,S:N211- | ON897715 |
| Conv-21 | BA.1 | S:A67V,S:T95I,S:Y145D,S:L212I,S:V320I,S:G339D,S:S371L,S:S373P,S:S375F,S:K417N,S:N440K,S:G446S,S:S477N,S:T478K,S:E484A,S:Q493R,S:G496S,S:Q498R,S:N501Y,S:Y505H,S:T547K,S:D614G,S:Q628K,S:H655Y,S:N679K,S:P681H,S:N764K,S:D796Y,S:N856K,S:Q954H,S:N969K,S:L981F | S:H69-,S:V70-,S:G142-,S:V143-,S:Y144-,S:N211- | ON897722 |
| Conv-23 | B.1 | S:D614G | | OM000280 |
| Conv-24 | B.1 | S:D614G | | OM000281 |
| Conv-25 | B.1 | S:D614G | | OM000284 |
| Conv-26 | B.1 | S:D614G | | OM000285 |
| Conv-27 | B.1 | S:D614G | | OM000282 |
| Conv-28 | B.1.2 | S:D614G | | OM000296 |
| Conv-29 | B.1 | S:D614G | | OM000286 |
| Conv-30 | B.1 | S:D614G | | OM000279 |
| Conv-31 | B.1 | S:D614G | | OM000288 |
| Conv-32 | B.1 | S:D614G | | OM000287 |
| Conv-33 | B.1 | S:D614G | | OM000283 |
| Conv-34 | B.1.2 | S:D614G | | OM000297 |
| Conv-35 | B.1.2 | S:D614G | | OM000299 |
| Conv-36 | B.1.2 | S:D614G | | OM000298 |
| Conv-37 | B.1.2 | S:D614G | | OM000294 |
| Conv-38 | B.1.1.207 | S:D614G,S:P681H | | ON897726 |
| Conv-39 | B.1.1.207 | S:E484K,S:D614G,S:P681H | | ON897727 |
| Conv-40 | B.1.1.207 | S:E484K,S:D614G,S:P681H | | ON897728 |
| Conv-41 | B.1.2 | S:G257D,S:D614G | | OM000295 |
| Conv-42 | P.1.10 | S:L18F,S:T20N,S:P26S,S:D138Y,S:R190S,S:K417T,S:E484K,S:N501Y,S:D614G,S:H655Y,S:S704L,S:T1027I,S:A1078S,S:V1176F | | ON897729 |
| Conv-43 | B.1.1.519 | S:L5F,S:T478K,S:D614G,S:P681H,S:T732A | | ON897730 |
| Conv-44 | B.1.526 | S:L5F,S:T95I,S:D253G,S:E484K,S:D614G,S:A701V | | ON897731 |
| Conv-45 | B.1.1.7 | S:N501Y,S:A570D,S:D614G,S:P681H,S:T716I,S:Q836*,S:S982A,S:D1118H | S:H69-,S:V70-,S:Y144-,S:I870-,S:A871-,S:Q872-,S:Y873-,S:T874-,S:S875-,S:A876-,S:L877-,S:L878-,S:A879-,S:G880-,S:T881-,S:I882-,S:T883-,S:S884-,S:G885-,S:W886-,S:T887-,S:F888-,S:G889-,S:A890-,S:G891-,S:A892-,S:A893-,S:L894-,S:Q895-,S:I896-,S:P897-,S:F898-,S:A899-,S:M900-,S:Q901-,S:M902-,S:A903-,S:Y904-,S:R905-,S:F906-,S:N907-,S:G908-,S:I909-,S:G910-,S:V911-,S:T912-,S:Q913-,S:N914-,S:V915-,S:L916-,S:Y917-,S:E918-,S:N919-,S:Q920-,S:K921-,S:L922- | OM000289 |
| Conv-46 | B.1.1.7 | S:N501Y,S:A570D,S:D614G,S:P681H,S:T716I,S:S982A,S:D1118H | S:H69-,S:V70-,S:Y144- | OM000292 |
| Conv-47 | B.1.1.7 | S:N501Y,S:A570D,S:D614G,S:P681H,S:T716I,S:S982A,S:D1118H,S:K1191N | S:H69-,S:V70-,S:Y144- | OM000291 |
| Conv-48 | B.1.429 | S:S13I,S:T95I,S:W152C,S:L452R,S:D614G | | ON897732 |
| Conv-49 | B.1.429 | S:S13I,S:W152C,S:L452R,S:D614G | | ON897733 |
| Conv-51 | AY.74 | S:T19R,S:G142D,S:R158G,S:A222V,S:L452R,S:T478K,S:D614G,S:P681R,S:D950N | S:E156-,S:F157- | OM000262 |
| Conv-52 | AY.62 | S:T19R,S:G142D,S:R158G,S:A222V,S:L452R,S:T478K,S:D614G,S:P681R,S:G946V,S:D950N | S:E156-,S:F157- | OM000270 |
| Conv-53 | AY.47 | S:T19R,S:G142D,S:R158G,S:A222V,S:V289I,S:L452R,S:T478K,S:D614G,S:P681R,S:D950N | S:E156-,S:F157- | OM000264 |
| Conv-54 | AY.25.1 | S:T19R,S:G142D,S:R158G,S:L452R,S:T478K,S:D614G,S:P681R,S:D950N | S:E156-,S:F157- | OM000271 |
| Conv-55 | AY.14 | S:T19R,S:G142D,S:R158G,S:L452R,S:T478K,S:D614G,S:P681R,S:D950N | S:E156-,S:F157- | OM000267 |
| Conv-56 | AY.14 | S:T19R,S:G142D,S:R158G,S:L452R,S:T478K,S:D614G,S:P681R,S:D950N | S:E156-,S:F157- | OM000266 |
| Conv-57 | AY.3 | S:T19R,S:G142D,S:R158G,S:L452R,S:T478K,S:D614G,S:P681R,S:D950N | S:E156-,S:F157- | OM000274 |
| Conv-59 | B.1.617.2 | S:T19R,S:K77T,S:G142D,S:R158G,S:G181V,S:L452R,S:T478K,S:D614G,S:A653V,S:P681R,S:D950N | S:E156-,S:F157- | OM000265 |
| Conv-60 | B.1.617.2 | S:T19R,S:K77T,S:G142D,S:R158G,S:G181V,S:L452R,S:T478K,S:D614G,S:A653V,S:P681R,S:D950N | S:E156-,S:F157- | OM311576 |
| Conv-61 | AY.25 | S:T19R,S:S112L,S:G142D,S:R158G,S:L452R,S:T478K,S:D614G,S:P681R,S:D950N | S:E156-,S:F157- | OM000263 |
| Conv-62 | AY.25 | S:T19R,S:S112L,S:G142D,S:R158G,S:L452R,S:T478K,S:D614G,S:P681R,S:D950N | S:E156-,S:F157- | OM000268 |
| Conv-63 | AY.44 | S:T19R,S:T22I,S:G142D,S:R158G,S:L452R,S:T478K,S:D614G,S:P681R,S:D950N | S:E156-,S:F157- | OM000272 |
| Conv-64 | AY.100 | S:T19R,S:T95I,S:G142D,S:R158G,S:L452R,S:T478K,S:D614G,S:P681R,S:D950N | S:E156-,S:F157- | OM000276 |
| Conv-65 | AY.119 | S:T19R,S:T95I,S:G142D,S:R158G,S:L452R,S:T478K,S:D614G,S:P681R,S:D950N | S:E156-,S:F157- | OM000273 |
| Conv-66 | AY.100 | S:T19R,S:T95I,S:G142D,S:R158G,S:L452R,S:T478K,S:D614G,S:P681R,S:D950N,S:G1124V | S:E156-,S:F157- | OM000269 |

**Table S8. SARS-CoV-2 variant spikes of human primary infection serum samples commercially obtained**

| Specimen ID | Infecting Genotype | Spike mutations |
|---|---|---|
| 738741 | B.1 | S:S13I;S:Q52R;S:A67V;S:L452R |
| 743259 | B.1 | D614G |
| 718055 | B.1.2 | D614G |
| 718057 | B.1.2 | S24L;S:D614G |
| 743136 | B.1.234 | D614G |
| 743256 | B.1.429 | S:W152C;S:L452R;S:D614G |
| 743257 | B.1.429 | S:L452R;S:D614G |
| 743264 | B.1.429 | S:S13I;S:W152C;S:L452R;S:D614G |
| 719166 | B.1.577 | D614G |
| 743255 | C.11 | S:L452R;S:D614G |
| D000113656 | B.1.1.7 | S:N501Y;S:A570D;S:D614G;S:P681H;S:T716I;S:S982A;S:D1118H |
| D000113657 | B.1.1.7 | S:N501Y;S:A570D;S:D614G;S:P681H;S:T716I;S:S982A;S:D1118H |
| D000113667 | B.1.1.7 | S:V433F;S:A570D |
| D000113669 | B.1.1.7 | S:N501Y;S:A570D;S:D614G;S:P681H;S:T716I;S:S982A;S:D1118H |
| D000113675 | B.1.1.7 | S:V193L;S:W436*;S:V510L;S:A1020S;S:P1079S;S:D1118H |
| D000113694 | B.1.1.7 | S:N501Y;S:A570D;S:D614G;S:P681H;S:T716I;S:S982A;S:D1118H;S:P1263L |
| D000117099 | C.37 | S:G75V;S:T76I;S:R246N;S:L452Q;S:F490S;S:D614G;S:T859N |
| D000117104 | C.37 | S:G75V;S:T76I;S:R246N;S:L452Q;S:F490S;S:D614G;S:T859N |
| D000117136 | C.37 | S:G75V;S:T76I;S:R246N;S:D442Y;S:L452Q;S:F490S;S:Q580X;S:T581X;S:D614G;S:I714V |
| D000117154 | C.37 | S:G75V;S:T76I;S:R246N;S:L452Q;S:F490S;S:D614G;S:T859N |
| D00011366 | B.1.1.7 | S:N501Y;S:A570D;S:P681H;S:T716I;S:S982A;S:D1118H |
| D00012307 | P.1 | S:E484K;S:N501Y;S:D614G;S:H655Y;S:Q677R;S:T1027I;S:V1176F |
| D00012308 | C.37 | S:G75V;S:T76I;S:R246N; S:S247-;S:Y248-;S:L249-;S:T250-;S:P251-;S:G252-;S:D253-;S:L452Q;S:D614G;S:Q675H;S:I720V;S:T859N;S:P863L;S:Q1180 |
| D00012308 | C.37 | S:G75V;S:T76I;S:R246N; S:S247-;S:Y248-;S:L249-;S:T250-;S:P251-;S:G252-;S:D253-;S:L452Q;S:F490S;S:D614G;S:I714V;S:T859N;S:G1219C |
| D00012308 | C.37 | S:W64-;S:H66-;S:A67-;S:I68-; S:T63X;S:F65X;S:H69-;S:V70-;S:S71-;S:G72-;S:T73-;S:N74-;S:G75-;S:T76-; S:D138Y;S:R246N;S:S247-;S:Y248-;S:L249-;S:T250-;S:P251-;S:G252-;S:D253- S:L452Q;S:F490S;S:D614G;S:T859N |
| D000123091 | C.37 | S:G75V;S:T76I; S:S247-;S:Y248-;S:L249-;S:T250-;S:P251-;S:G252-;S:D253-S:R246N;S:L452Q;S:D614G;S:T859N |
| D000123095 | C.37 | S:G75V;S:T76I; S:S247-;S:Y248-;S:L249-;S:T250-;S:P251-;S:G252-;S:D253-;S:R246N;S:L452Q;S:F490S;S:D614G;S:T859N |
| D000123099 | C.37 | ;S:G75V;S:T76I; S:S247-;S:Y248-;S:L249-;S:T250-;S:P251-;S:G252-;S:D253-;S:R246N;S:L452Q;S:F490S;S:D614G;S:T859N |
| D000123108 | P.1 | S:D138Y;S:R190S;S:K417T;S:E484K;S:N501Y;S:D614G;S:H655Y;S:Q677R;S:T1027I;S:V1176F |
| D000123182 | P.1 | S:L18F;S:T20N;S:P26S;S:D138Y;S:R190S;S:K417T;S:E484K;S:N501Y;S:D614G;S:H655Y;S:T1027I;S:V1176F |
| D000123194 | P.1 | S:D138Y;S:K417T;S:E484K;S:N501Y;S:D614G;S:H655Y;S:T1027I;S:V1176F |
